# Primate anterior insular cortex represents economic decision variables postulated by Prospect theory

**DOI:** 10.1101/2020.11.12.380758

**Authors:** You-Ping Yang, Xinjian Li, Veit Stuphorn

## Abstract

In humans, risk attitude is highly context-dependent, varying with wealth levels or for different potential outcomes, such as gains or losses. These behavioral effects are well described by Prospect Theory, with the key assumption that humans represent the value of each available option asymmetrically as gain or loss relative to a reference point. However, it remains unknown how these computations are implemented at the neuronal level. Using a new token gambling task, we found that macaques, like humans, change their risk attitude across wealth levels and gain/loss contexts. Neurons in their anterior insular cortex (AIC) encode the ‘reference point’ (i.e. the current wealth level of the monkey) and the ‘asymmetric value function’ (i.e. option value signals are more sensitive to change in the loss than in the gain context) as postulated by Prospect Theory. In addition, changes in the activity of a subgroup of AIC neurons are correlated with the inter-trial fluctuations in choice and risk attitude. Taken together, we find that the role of primate AIC in risky decision-making is to monitor contextual information used to guide the animal’s willingness to accept risk.

## Introduction

Uncertainty about the possible outcomes of chosen actions is a basic feature of all human and animal decision making. How our nervous system deals with this uncertainty is therefore a fundamental question in cognitive neuroscience. Decisions under uncertainty depend on an individual’s risk attitude, i.e., the willingness to accept uncertainty about the outcome (risk) in exchange for possibly better outcomes than a safer alternative. Risk attitude is strongly influenced by context. Humans show different risk attitudes when facing risky gains versus risky losses^1^. The abundance of economic resources in the environment and the current wealth of subjects also modulate an individual’s risk attitude^2–5^. Prospect theory^6^, the most influential^7^ and wide-ranging^8^ descriptive model of decision-making under risk, explains these context-dependent changes in risk attitude using two critical concepts about the cognitive processes underlying value estimation. First, prospect theory assumes that humans evaluate possible future outcomes either as gains or as losses relative to a reference point (i.e. the current wealth, resources, or state of the subject). Second, human’s sensitivities to changes in value are different for losses and gains. Specifically, humans are more sensitive to the change of the value of a loss as compared to a gain (i.e. *losses loom larger than gains)*. Thus, humans’ choices can be manipulated by framing an identical outcome as either a gain or loss using verbal instruction, and by varying the current wealth of subjects that change the point of reference. Despite the success that the prospect theory has achieved in explaining the risky choices of humans, it remains unclear how the value function across (gain/loss) contexts, as well as the point of reference are represented in our brain on the neuronal level.

Human imaging experiments and lesion studies have identified a network of brain areas that are active during decision-making under risk^9–14^. Of particular interest is the anterior insular cortex (AIC), a large heterogeneous cortex in the depth of the Sylvian fissure. Human fMRI studies have suggested a crucial role of AIC in representing subjects’ current internal states^15,16^, and in risk-aversive behavior^12,13^. Lesions in the AIC have also been documented to affect the risk-attitude of human patients^17,18^. Moreover, recording studies in monkeys have shown that AIC neurons encode reward expectation^19,20^. Based on these findings, we hypothesized that the AIC neurons may encode behaviorally relevant contextual information in the framework suggested by prospect theory. AIC would represent the current state of the subject (the reference point) as well as reference-dependent value signals that differ in loss or gain context (asymmetrical value functions in loss and gain). Together, these representations in AIC would influence a subject’s risk attitude in decision making.

To test this hypothesis, we developed a token-based gambling task and recorded single neuron activities from the AIC of two macaque monkeys engaged in this task. We first examined whether and how monkeys changed their risk attitude in various behavioral contexts. Next, we identified AIC neurons representing factors that influence risk attitude, such as starting token number, gain or loss outcome, total value of the option, and uncertainty. Finally, we determined whether the AIC neurons encoding these factors also predict the monkey’s choice or risk attitude.

We found that monkeys, like humans, have different risk attitudes depending on the gain/loss context, and that AIC neurons encode reference-dependent value signals, consistent with the asymmetric value function as postulated by the Prospect theory. In addition, both the monkeys’ choices and the activity of AIC neurons are strongly influenced by the number of tokens that the monkeys possessed at the start of the trial, indicating that the momentary wealth level served as a reference point. Inter-trial fluctuations in the activity of AIC neurons encoding these variables were correlated with the monkeys’ choices and risk attitude. Taken together, these results support our hypothesis that the primate AIC encodes the reference point and reference-dependent value signals, and that these value representations of available options modulate the animal’s willingness to accept risk in the current behavioral context.

## Result

Two monkeys were trained in a token-based gambling task (**Figure 1a**). In this task, the monkey had to collect a sufficient number of tokens (≥6) to receive a standard fluid reward (600μl water). Because the maximum number of tokens that could be earned in a single trial was three, the monkeys had to accumulate the necessary tokens over multiple trials (**Figure S1**). On each choice trial, the monkey chose between a gamble option (uncertain outcome) and a sure option (certain outcome), which could result in gaining or losing tokens. The number of tokens to be won or lost was indicated by the color of the target cues, while the probability was indicated by the relative proportion of each colored area (**Figure 1b**). To investigate whether the monkeys’ risk attitude was different for gains and losses, we presented either only gain or only loss options on any given trial. Thus, in a gain context, the monkey had to choose between a certain token increase versus an uncertain option that could result in an even larger increase or no increase at all (**Figure 1b, left**). In the loss context, the monkey had to choose between a certain loss and an uncertain option that could result in no loss at all or an even larger loss (**Figure 1b, right**). The monkeys selected the chosen option by making a saccade to the corresponding target cue. After a short delay (450-550ms), the outcome was revealed, and the number of currently owned tokens (token assets) was updated. If a trial ended with a token number less than 6 (e.g., 4), these tokens (e.g., 4) were kept as the start tokens for the next trial. If a trial ended with a token number larger than 6 (e.g., 8), beside a delivery of water, the remaining tokens (e.g., 8-6=2) were rolled over to the start of the next trial.

**Figure 1.**
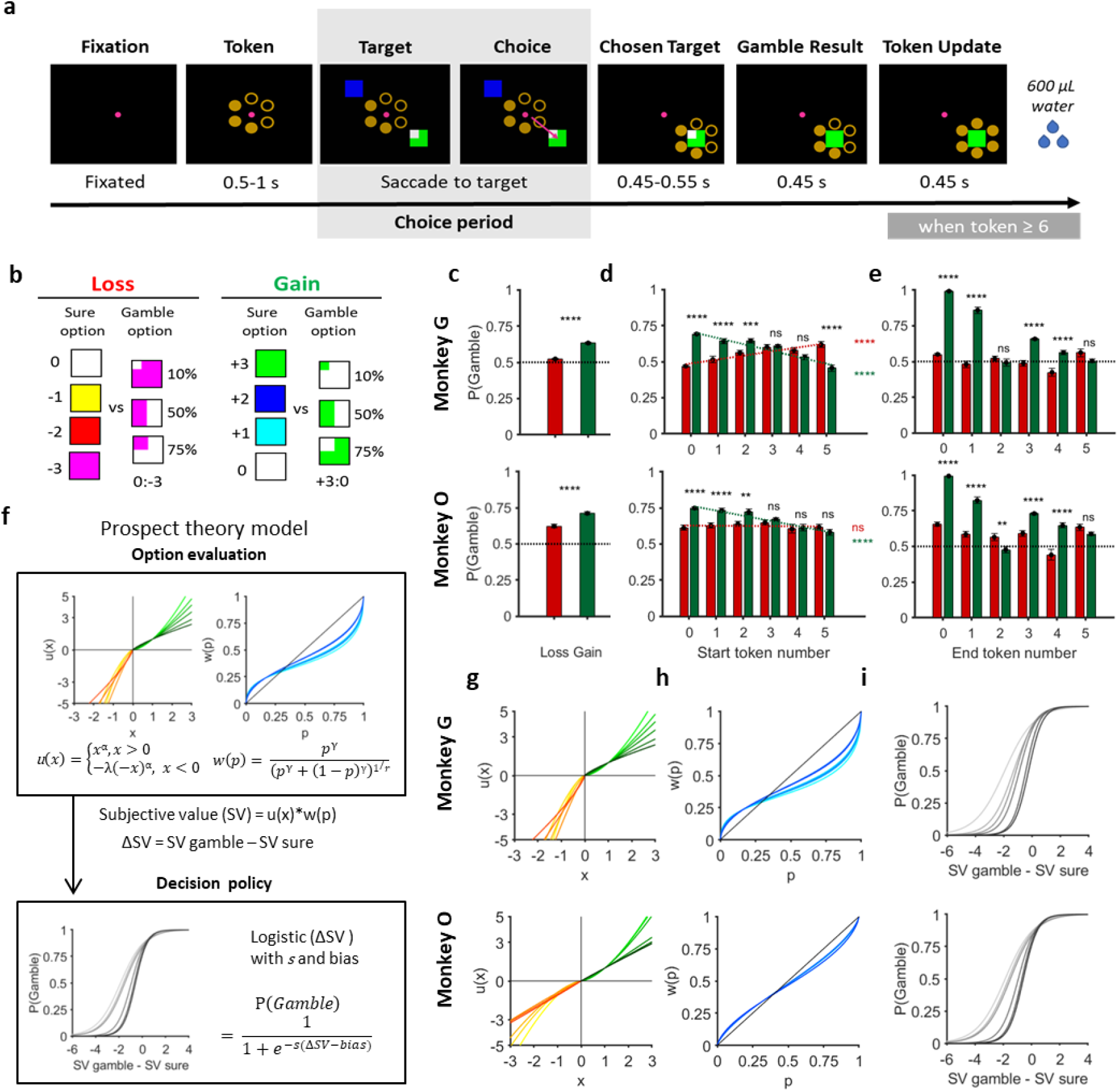
Behavior performance of monkeys in the token-based gambling task. (a) Schematic of the task. Each trial starts with a fixation dot at the center of the screen. Upon the monkey fixated to the central dot, the current number of tokens it has was presented (filled circles of the hexagonal placeholder). Following 0.5-1s delay, one (‘forced-choice’ trial) or two (‘choice’ trial) options were presented (detailed in (b)), and the monkey indicated its choice by making a saccade to the target. The unchosen option then disappeared, and the current number of tokens was presented again in the surround of the chosen target. The outcome of the chosen target revealed after a delay (0.45-0.55 s), indicated by the color change of the square, and the number of tokens that the monkey possessed was updated accordingly. The monkey was rewarded (600uL of water) whenever it collected six tokens or more at the end of the trial. Shadowed area indicated the choice period of which the neuronal data was analyzed. (b) Set of options. Each option is a square (x degree), of which the color(s) indicated the possible outcome(s) (−3 to +3, in units of token change), and the portion of colored area indicated the probability (10%, 50%, 75%, 100%) of the corresponding outcome to be realized. The choice trials consisted of two types (gain vs. loss context), and there was always a sure option paired with a gamble option – i.e., only the combination of [sure gain vs. gamble gain] and [sure loss vs. gamble loss] were available. In forced-choice trials, only one option was presented, which could be either a sure option or gamble option (gain or loss). See Methods for details. (c) The probability of monkey choosing gamble option in gain/loss contexts. Upper and lower panels represent data from two different monkeys, respectively. Green: gain context; Red: loss context. Error bars: S.E.M; **** p<10^−4^, paired t-test. (d) The probability of monkey choosing gamble option, plotted by context and the start token number. Green: gain context; Red: loss context. Error bars: S.E.M; n.s.: not statistically significant (p>0.05), ** p<10^−2^, *** p<10^−3^, **** p<10^−4^ in paired t-test (black), **** p<10^−4^ in regression analyses (green or red). (e) The probability of monkey choosing gamble option, plotted by context and the end token number. Conventions as in (d). (f) Behavior modeling. The model consists of two parts: first in the process of option evaluation (upper panel), the subjective value (SV) of each option was calculated as the product of a utility function and a probability weighting function. Both functions are nonlinear as per the Prospect theory (PT) hypothesized. The subjective value difference between the two options (ΔSV) was then used to determine the probability of choosing the gamble option via a logistic function –i.e., decision policy (lower panel). (g-i) The best fit utility functions (g), probability weighting functions (h), and the decision policies (i) based on the observed performance. Upper and lower panels represent data from two different monkeys, respectively. Color gradients represent different start token numbers (light to dark: 0 to 5).

Both monkeys learned the task, as indicated by the observation that their fixation behavior was strongly influenced by their token assets. Monkeys fixated faster (**Figure S2a-c**) and were less likely to break their fixation (resulting in abortion of the trial) (**Figure S2d-f**) when they had larger token assets at the start of the trial, and when they received more tokens from the previous trial. These results suggest that monkeys understood the use of tokens as secondary reinforcers, and thus were more motivated when they owned more and received more tokens, before they actually earned the primary reinforcer (the fluid reward).

### Monkeys’ risky choices are influenced by gain/loss context and current token assets

We found that monkeys’ choices were influenced by the gain/loss context. Both monkeys were more likely to choose the gamble option than the sure option (**Figure 1c**; t-test; Monkey G, P(Gamble)=59%, p<10^−4^; Monkey O, P(Gamble)=67%, p<10^−4^) and were even more likely to do so in the gain context than in the loss context (**Figure 1c**; paired t-test, p<10^−4^ for both Monkey G and Monkey O). We have also found that monkeys’ choices were influenced by the number of tokens they owned at the start of the trial (‘current token assets’), but differently for gains and losses. In the gain context, the probability of the monkey choosing the gamble option (P(Gamble)) decreased as the token assets increased (**Figure 1d**; green dashed line; regression analysis; Monkey G, β = −0.044, p<10^−4^; Monkey O, β = −0.035, p<10^−4^). In contrast, in the loss context P(Gamble) increased with increasing token assets (**Figure 1d**; red dashed line; regression analysis; Monkey G, β = 0.028, p<10^−4^; Monkey O, β = −0.001, p = 0.8). Thus, as the monkeys owned more token assets, they became more risk-averse for further gains (i.e., less willing to gamble for a greater win), but were more risk-seeking for avoiding a potential loss. These results are in line with the observation of humans that human subjects tend to be more risk-aversive when facing a potential gain, and more risk-seeking when facing a potential loss as their own asset increases^1^.

Monkeys’ response times (RTs, the interval between stimulus onset and the saccade initiation) were also influenced by these contextual factors. Both monkeys responded slower in the loss context than in the gain context (Figure S3a-b; permutation test; monkey G: RT _gain_ = 204.79ms, RT_loss_ = 246.51ms, p < 10^−3^; monkey O: RT _gain_ = 175.07ms, RT_loss_ = 206.00ms, p < 10^−3^), and when they had more tokens (**Figure S3c-d**; regression analysis; monkey G: β_StartTkn_ = 2.83, p = 0.19; monkey O: β_StartTkn_ = 3.50, p < 10^−2^). These suggest that monkeys chose more carefully when facing a potential loss, and when they are getting closer to 6 tokens for the water reward. Other factors that impacted RTs will be discussed in later sections (**Figure S3e-h**).

The fact that monkeys’ risk preference changed across contexts suggests that they evaluate each available option not depending on the final status (the final token number), but rather how the current status will be changed (by gaining or losing tokens). Thus, the monkeys showed a framing effect^21^. This is clearly demonstrated by contrasting trials with the same outcome in terms of final token number, but which resulted from either gaining or losing tokens (**Figure 1e**; paired t-test; monkey G & O: p<10^−2^ for all end token numbers, except end token numbers 2&5 for monkey G, and number 5 for monkey O). Monkeys chose more gambles when a given prospect (end state) was offered as a gain, compared to when it was offered as a loss.

### Prospect theory model of risk-attitude adjustment

After confirming that monkeys’ choice behavior was influenced by core contextual factors critical for prospect theory (PT), we used this model to describe choice behavior (**Figure 1**). The key component of the PT model is to weight gains, losses, and probabilities differently before they are combined to form a subjective evaluation of the option. The relative gains and losses are mapped onto corresponding subjective utility as follows: u(x) = x^α^ when x>0 (reward outcome in gain) and u(x) = −λ*(-x)^α^ when x<0 (reward outcome in loss). The utility function component α captures risk-attitude. Convex utility (with α > 1) indicates risk-seeking, while subjects are more sensitive to differences in larger rewards. Concave utility (with α < 1) indicates risk-aversive, while subjects exhibit diminishing marginal sensitivity. The utility function component λ captures loss-aversion, the idea that losses loom larger than equivalent gains. λ > 1 indicates more sensitivity to losses than gains and λ < 1 indicates more sensitivity to gains than losses.

To capture the influence of tokens on different components, we modeled behavior for each start token number independently. In the gain context, both monkeys were risk-seeking (α > 1) when the start token number was low, but they became risk neutral or risk-averse when the start token number increased (**Figure 1g**; light to dark green lines indicate increasing start token number). Estimated α was negatively modulated by the start token number (regression analysis; Monkey G, β = −0.16, p<10^−4^; Monkey O, β = −0.14, p<10^−4^). In monkey G, the utility functions were consistently steeper for losses than for gains (**Figure 1g**; yellowish to red lines indicate increasing start token number; t-test: λ > 1; Monkey G, p<10^−4^ for all start token numbers). Thus, monkey G showed loss-aversion. However, in monkey O the utility functions were not consistently steeper for losses than for gains and λ values varied around 1. This indicated that monkey O was equally sensitive to gains and losses and thus showed no loss aversion. There was no significant difference for the estimated λ across different start token numbers for either monkey (regression analysis; Monkey G, β = 0.03, p = 0.69; Monkey O, β = 0.01, p=0.4).

Objective probabilities are mapped onto a subjective weighting function as follows: 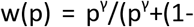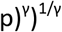. γ > 1 indicates an S-shape subjective probability mapping (overestimated for large probabilities and underestimated for small probabilities), γ < 1 indicates an inverse S-shape subjective probability mapping (underestimated for large probabilities and overestimated for small probabilities), and γ = 1 indicate a linear mapping of objective probabilities. Both monkeys showed an inverse S-shaped mapping of probabilities (**Figure 1h**; t-test: γ < 1; both monkeys, p<10^−4^ for all start token numbers). The mappings were slightly influenced by increasing start token numbers in one monkey (light blue to dark blue lines; regression analysis; Monkey G, β = 0.02, p<10^−4^) and not at all in the other one (Monkey O, β = 0.0004, p = 0.89).

After calculating the subjective value (SV = u(x)*w(p)) of each option based on prospect theory, we estimated the P(Gamble) by passing subjective value difference between options (ΔSV) through a softmax function with parameters s and bias: P(Gamble) = 1/1+e^−s(ΔSV-bias)^, where *s* controls choice stochasticity and *bias* represents the tendency to choose the gamble option independent of the value calculation process. The monkeys showed a consistent tendency to choose the gamble option (**Figure 1i**; leftward shift of the choice function in; t-test: both monkeys, p<10^−4^ for all start token numbers). This tendency decreased when the start token number increased (**Figure 1i**; light gray to black lines; regression analysis; Monkey G, β = 0.32, p<10^−4^; Monkey O, β = 0.28, p<10^−4^), indicating monkeys became less risk-seeking as their wealth levels increased. Moreover, the choice functions became steeper, that is monkeys’ choices became less stochastic, when the start token number increased for both monkeys (regression analysis; Monkey G, β = 0.23, p<10^−4^; Monkey O, β = 0.22, p<10^−4^). This result, combined with the token effect on response time, indicates that choices became slower but less stochastic when token assets increased, which suggests a speed-accuracy tradeoff.

In sum, the PT model describes the behavioral result well (**Figure 1c-d**) and predicts the monkeys’ choices better than the expected value model that does not assume nonlinearity in subjective utility and probabilities (**Figure S3**). This was also true after taking into account the different number of free parameters (see quantitative model evaluation in **Table 1**). This suggests that key components of the prospect theory model are important for explaining the monkeys’ behavior in our task.

**Table 1.**
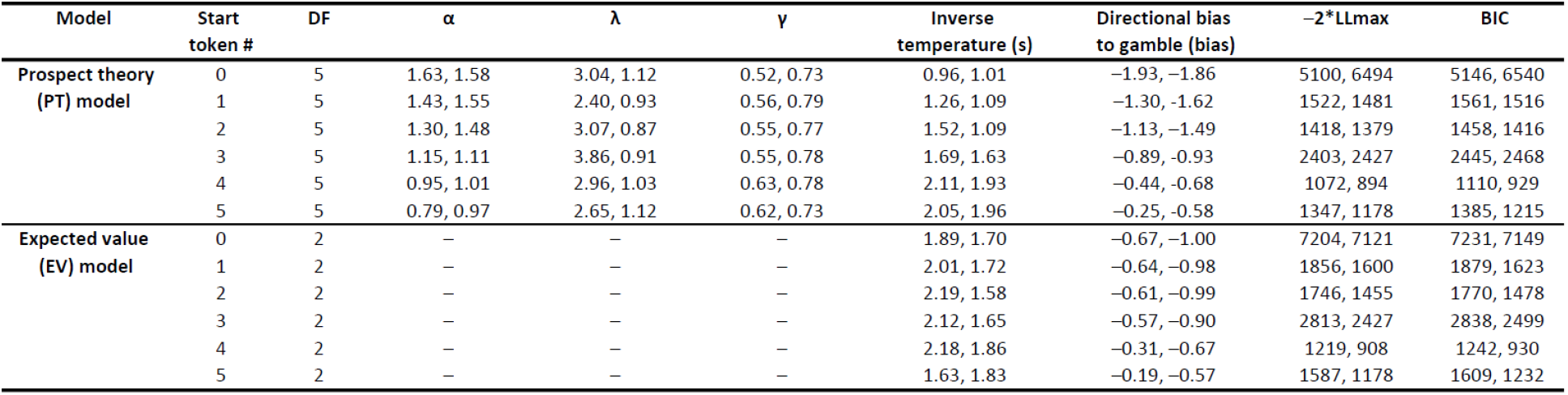
Model comparison. DF, degrees of freedom; LLmax, maximal log likelihood; BIC, Bayesian Information Criterion (computed with LLmax). The table summarizes for each model the likelihood maximizing (‘best’) parameters average across sessions and its fitting performances for each monkey. Comparing the model fit of PT model and EV model: t-test; Monkey G, p < 10^−4^, p<10^−4^, p<0.05, p<0.05, p=0.09, and p<0.05 for start token number 0-5, respectively; Monkey O, p <10^−2^, p<10^−3^, p=0.55, p= 0.62, p=0.95, and p=0.80 for start token number 0-5, respectively. Comparing the BIC of PT model and EV model: t-test; Monkey G, p <10^−4^, p<10^−4^, p<10^−2^, p<0.05, p<0.05, and p<0.01; Monkey O, p <10^−2^, p<10^−4^, p=0.22, p=0.17, p=0.20, and p=0.34 for start token number 0-5, respectively.

### Anterior Insula neurons encode decision-related variables that influence risk-attitude

To determine the neuronal basis underlying prospect theory, we recorded 240 neurons in the AIC of two macaque monkeys (monkey G: 142 neurons; monkey O: 98 neurons) working in the token gambling task. The recording locations are shown in **Figure 2a** (more details in **Figure S5)**. We analyzed the neuronal activity in the choice period (i.e., the time from target onset to saccade initiation) to determine if AIC neurons carried signals that could influence decision making. We began by analyzing activity during no-choice trials, in which only one option was presented. In general, the AIC neurons showed weak spatial selectivity. Only 7% (17/240) of all AIC neurons showed a significant effect of spatial location on neuronal activity (1-way ANOVA, p<0.05). We therefore ignored spatial target configuration for the remaining analysis.

**Figure 2.**
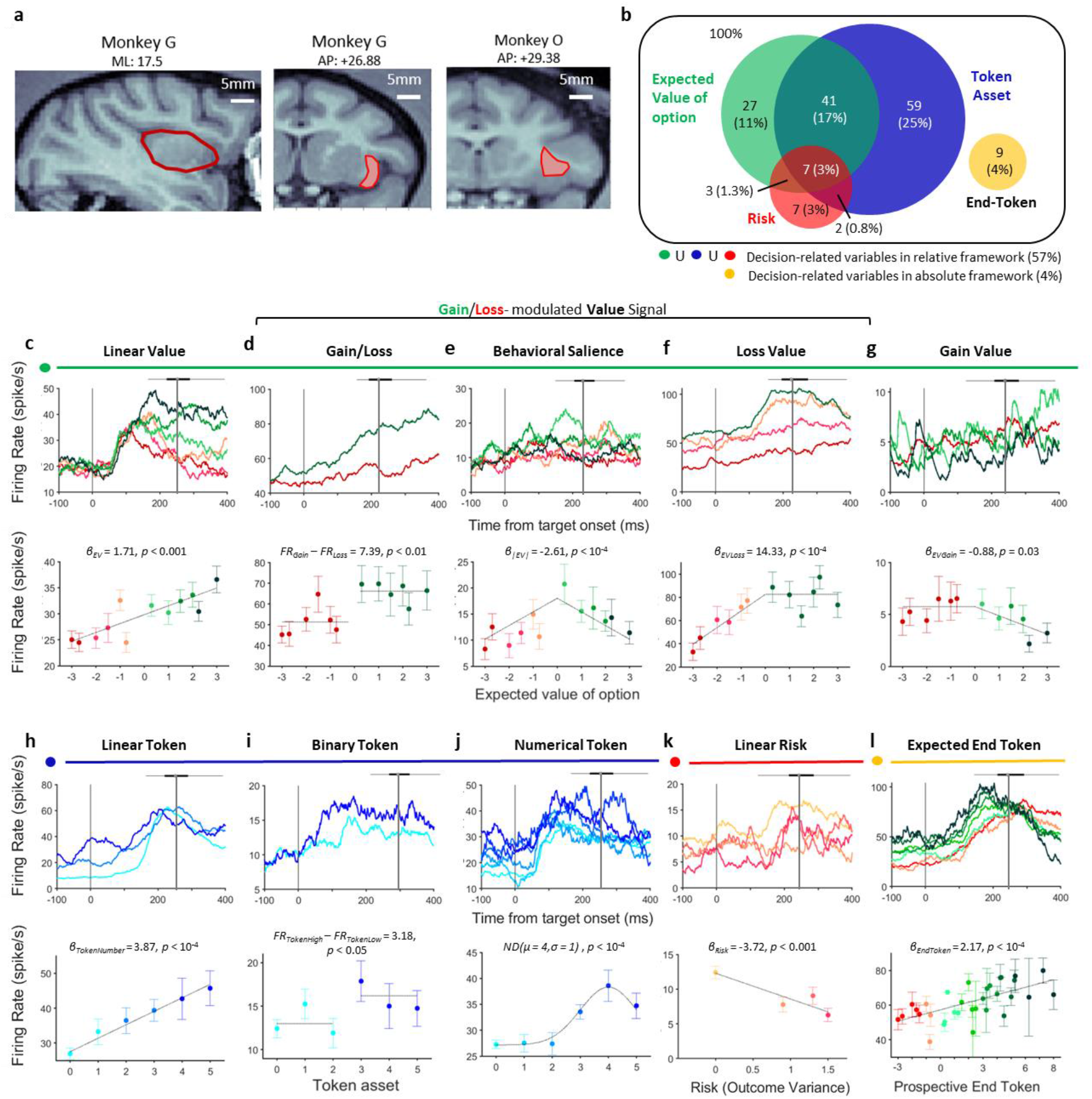
AIC Neurons encode diverse task-related variables in forced-choice trials. (a) MRI images showing the area of recording of each monkey. Left and middle: sagittal (left) and coronal (middle) view of the insular cortex of monkey G. Right: coronal view of the insular cortex of monkey O. (b) Venn diagram of the neurons encoding four task-related variables in the forced-choice trials. Green: expected value of option (EV); Blue: start token number; Red: risk (variability of potential outcomes); Yellow: end token number. (c-g) Example neurons showing a variety of patterns by which the contextual information (gain vs. loss) and/or the EV were encoded. Upper panels: spike density function (SDF), aligned by the target onset (t=0). Lower panels: mean firing rate of each example neuron at different EV levels. Mean firing rate was calculated using the window from target onset to saccade initiation (varied across trials). The distribution of saccade timing was presented as a boxplot on top of each SDF. For clarity, when plotting the SDF, data of some EV levels were grouped together, as indicated by the color codes. (c) linear encoding of the EV across contexts; (d) binary encoding of gain/loss context; (e) linear encoding of the absolute value of the EV in both contexts; (f) linear encoding of the EV in the loss context; (g) linear encoding of the EV in the gain context. (h-j) Example neurons showing a variety of patterns by which the token information was encoded. (h) linear; (i) binary encoding of the start token number; (j) encoding of a specific number of start token (=4). Conventions are as in (c-g). (k) Example neuron showing linear encoding of the risk. Conventions are as in (c-g). (l) Example neuron showing linear encoding of the end token number. Conventions are as in (c-g).

To quantitatively characterize the variables that each AIC neuron encodes during the choice period, we examined the activity of each neuron using a series of linear regression models with all potential combinations of 3 basic variables (token assets, value, and risk) and a baseline term. This resulted in 8 families of models. Within each family, a basic variable could be represented by varying numbers of specific instantiations (3 forms of token-encoding, 5 forms of value-encoding, 2 forms of risk-encoding). This resulted in a total of 162 tested models, including a model that included only a baseline term (for details see Methods). For each neuron, we identified the best fitting model using the Akaike information criterion and classified it into different functional categories according to the variables that were most likely encoded by the neuronal activity.

The vast majority of recorded AIC neuron activity (146/240; 61%) encoded at least one decision-related variable (task-related neurons: p < 0.05 for the coefficient of a specific variable in the best-fitting multiple linear regression mode; **Figure 2b**, more details in **Table 1**). Among these task-related neurons, 78 neurons (33%) carried gain/loss-modulated value signals, 109 neurons (45%) carried token signals, 19 neurons (8%) carried risk signals, and 9 neurons (4%) carried an absolute value signal. A substantial number of AIC neurons (53/146; 36%) showed mixed selectivity and encoded more than one decision-related variable. The distributions of neural type classification were similar across the two monkeys (**Table S1**).

A large group of AIC neurons reflected information about the expected value of the options (value-encoding neurons). A subset of this group of AIC neurons, carrying a *Linear value signal* (example neuron in **Figure 2c**), encoded the expected value of all options in a monotonically rising (n=13) or falling (n=1) fashion, for both gains and losses. This kind of value signal is not gain/loss context sensitive. However, we found neuronal activity of other subsets of value-encoding neurons that varied largely as a function of the way they represented value across the Gain and the Loss context (**Figure 2d-e**). One group of these value-encoding neurons carried a binary *Gain/Loss signal* (**Figure 2d**) that categorized each option as gain or loss, regardless of the expected value. In addition, we found AIC neurons that represented value in both the gain and loss context, but with inverse correlations of neural activity and value (**Figure 2e**). These neurons likely carried a *Behavioral salience signal*. Most interestingly, we found two other groups of AIC neurons carrying *Loss value* (**Figure 2f**) or *Gain value signals* (**Figure 2g**), respectively. These neurons represented a value signal, but only in either the loss or the gain context. We encountered more Loss value neurons (n=29) than Gain value neurons (n=4). The larger number of neurons encoding Loss value fits with human neuroimaging findings that suggest a role for the anterior insula in encoding aversive stimuli and situations^1,13^.

The largest proportion of AIC neurons reflected information about the currently owned token number. These token-encoding neurons used three different frameworks for encoding token assets. The first group carried a *Linear token signal* (**Figure 2h**). These AIC neurons monotonically increased (n=11) or decreased (n=2) their activity with the number of token assets. The second group carried a *Binary token signal* (**Figure 2i**). These AIC neurons categorized all possible token numbers into a high [3, 4, 5] and a low [0, 1, 2] token level. Likely, this reflects a fundamental distinction between a ‘low’ token level, for which it is impossible that the monkey will earn reward at the end of the current trial (because the monkey can only earn a maximum of 3 tokens in one trial), and a ‘high’ token level that makes it possible to earn a reward in the current trial. The third, and largest, group carried a *Numerical token signal*. These AIC neurons are number-selective and are tuned for a preferred number (here four, example neuron in **Figure 2j**). We used a Gaussian function to fit this activity pattern. The AIC neurons carrying a Numerical token signal covered the entire scale from 0-5 tokens with some neurons having each of the possible token amounts as their preferred number.

Based on human neuroimaging data^22,23^, it has been suggested that the anterior insular cortex encodes the riskiness of options. We tested therefore if AIC neurons encoded risk-related signals. Here risk was defined as outcome variance. We found some AIC neurons (n=9) carrying a *Linear risk signal* (**Figure 2k**) that encoded the risk of the various options across both the gain and loss context in a parametric fashion. We also found another group of AIC neurons (n=10) encoding a *Binary risk signal* that categorized options into safe or uncertain. More details of distribution of type of neuron can be found in **Table 2** and **TableS2**.

**Table 2.** Summary of the number and percentage of significant responding neurons in different subsets of neuron types for all recorded AIC neurons.

|  |  |  |  |  |  |  |  |  |  |
| --- | --- | --- | --- | --- | --- | --- | --- | --- | --- |
| Gain/Loss-Value | Gain/Loss<br>13 (5%) | Loss Value<br>29 (12%) | Gain Value<br>4 (2%) | Behavioral Saliency<br>13 (5%) | Linear Value<br>14 (6%) |  |  |  | All<br>73 (30%) |
| Token | Linear Token<br>13 (5%) | Binary Token<br>11 (5%) | Token 0<br>11 (5%) | Token 1<br>16 (7%) | Token 2<br>24 (10%) | Token 3<br>12 (5%) | Token 4<br>10 (4%) | Token 5<br>8 (3%) | All<br>105 (44%) |
| Risk | Linear Risk<br>9 (4%) | Binary Risk<br>10 (4%) |  |  |  |  |  |  | All<br>19 (8%) |
| Expected End Token | End Token<br>9 (4%) |  |  |  |  |  |  |  | End Token<br>9 (4%) |

In the analysis so far, we have used a relative framework for value. Expected value was defined as token changes relative to a reference point (the start token number). However, value could also be defined in an absolute framework (i.e., the final token number at the end of the trial). Such an absolute value framework is arguably better suited for capturing the interest of the monkey in ascertaining how close he is to collecting 6 tokens and reaping a reward. We tested for AIC neurons that represented expected absolute value, which is the expected end token number weighted by the probability of each outcome. However, we found only a very small number (9/240; 4%) carrying an *End token signal* (**Figure 2l**).

A significant number of AIC neurons showed activity pattern that matched several predictions of prospect theory. First, we found that many AIC neurons encode the wealth level of the monkey, i.e. the token number at the start of the trial. Within the context of our task, this variable represented the reference point relative to which the gain or loss options are measured. Simultaneously, this variable also indicates the current state of progress and indicates how close the monkey is to achieving the next reward. Second, many other AIC neurons reflect in their activity whether the offer is a gain or a loss. Some of them encoded the context, i.e., whether the options were presented in the gain or loss context. Other neurons represented a gain/loss-specific value signal in a parametric manner exclusively. Third, only very few neurons encode expected absolute value. Taken together, these three findings strongly imply that the primate AIC uses a relative value encoding framework, anchored to a reference point that reflects the current state of the monkey, as suggested by prospect theory.

### Value-encoding neurons in AIC exhibited contextual modulation predicted by the Prospect theory

The majority of value-encoding AIC neurons were context-modulated (**Figure 2d-g**). A strong assumption of Prospect theory is that changes in relative value are not encoded symmetrically across gains and losses. Indeed, the monkeys’ behavior indicated that they were more sensitive to objective value differences in the loss than the gain context (i.e. steeper utility functions in the loss than that in gain context in **Figure 1g**). We therefore investigated whether and how value signals across the AIC population showed matching differences in their sensitivity for gains and losses. We examined the absolute value of the standardized regression coefficients (SRC) of Loss-Value Neurons in the loss context and that of Gain-Value Neuron in the gain context. At the population level, we found indeed that Loss value signals and Gain Value signals had different sensitivities to changes in value. Specifically, the normalized |SRC| of Loss-Value Neurons in the loss context were larger than that of Gain-Value Neuron in the gain context (**Figure 3a**, permutation test; mean of |SRC_loss_|= 2.978, mean of |SRC_gain_|= 2.058, p = 0.054; unsigned SRC for losses and gains were indicated in red and green, respectively). This suggests that the AIC neurons encoding value signals were more sensitive to increasing loss than increasing gain (**Figure 3b**).

**Figure 3.**
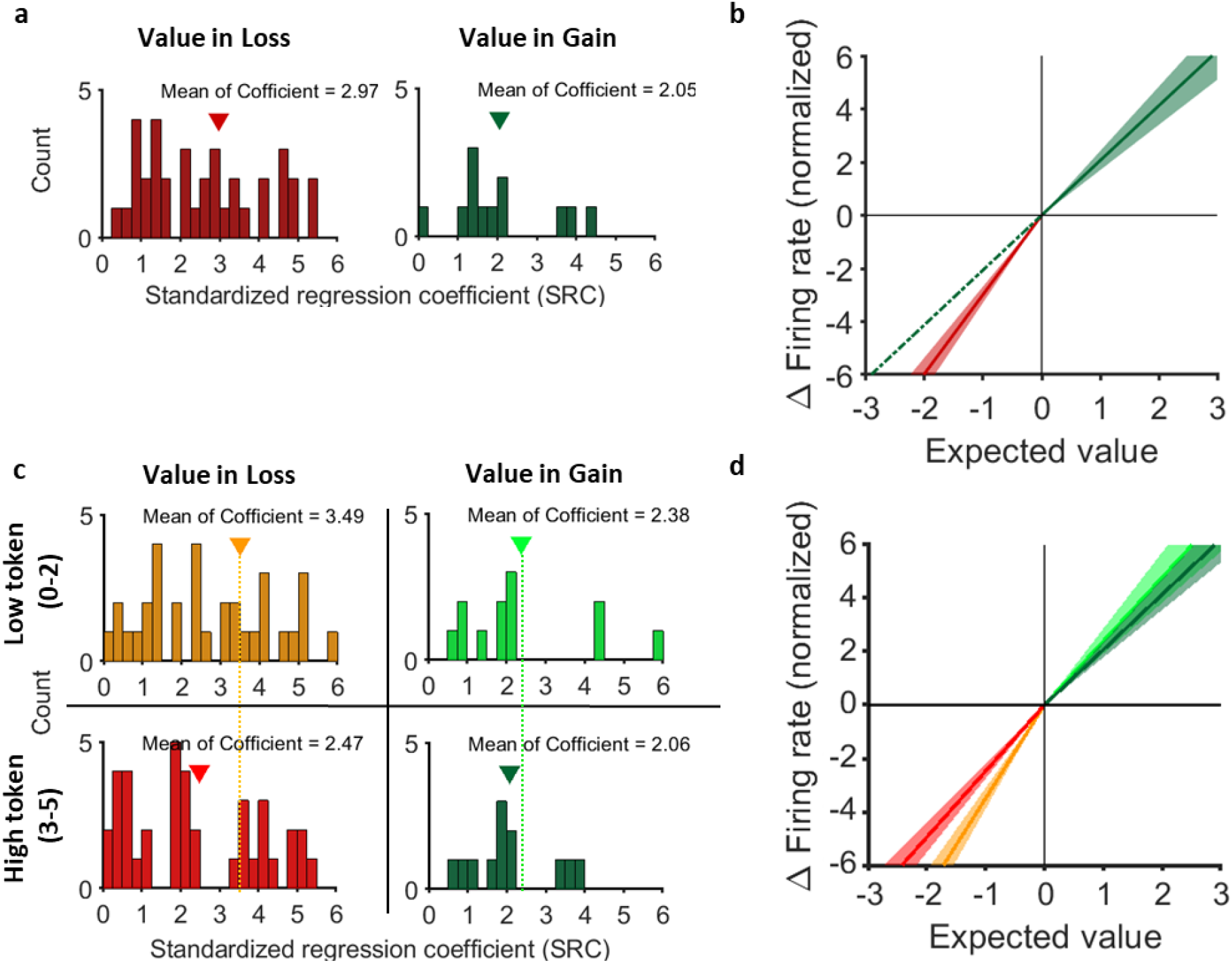
Gain-Value and loss-Value neurons exhibit differential sensitivity to EV change in the gain and loss context. (a-b) Linear regression was performed using the expected value of option (EV) as the regressor, to account for the variability of firing rates for Loss-Value neurons (39 neurons) in loss-context trials; and for Gain-Value neurons (12 neurons) in gain-context trials. See Methods for details of the neuron selection. (a) Distribution of the standardized regression coefficients (SRC). For cross-context comparison, the absolute value of SRCs (|SRCs|) were plotted. Left panel: data of those Loss-Value neurons in loss-context trials. Right panel: data of those Gain-Value neurons in gain-context trials. Each count represents one neuron. Inverted triangle: mean of the distribution. (b) Replot the SRCs (as the slope of ΔFR to ΔEV) of Loss Value neurons (red) and those of Gain Value neurons (green) in loss- and gain-context, respectively. Noted that the slope of red line is steeper than the slope of green line (p=0.053, permutation test), indicating that as compared to the Gain Value neurons, the Loss Value neurons are more sensitive to EV change (in the loss context), mirroring the pattern observed from behavior (Fig.1g). Solid line and shadow area: mean ± S.E.M. (c-d) Linear regression was performed using the expected value of option (EV) as the regressor, to account for the variability of firing rates for Loss-Value neurons in loss-context trials in low or high token level; and for Gain-Value neurons in gain-context trials in low or high token level. (c) Distribution of the |SRCs| of Loss Value neurons in loss-context (left column) and |SRCs| of Gain Value neurons in gain-context (right column), split by start token levels. Upper row: low token level (start token number = 0-2; Bottom row: high token level (start token number = 3-5). Conventions as in (a). (d) Replot the SRCs from (c). Note that for both gain- and loss-contexts, the slope becomes shallower as the token level increases, consistent to the pattern observed from behavior (Fig.1g).

Moreover, the sensitivity of value change in gain or loss context were also influenced by the wealth level. Normalized |SRC| of Loss-Value Neurons in the loss context became smaller as the wealth level increased (**Figure 3c**, left; permutation test; mean of |SRC_loss_| in low wealth level = 3.49, mean of |SRC_loss_| with high wealth level= 2.47, p = 0.017; unsigned SRC in the loss context for low or high wealth levels were indicated in orange and red, respectively). Normalized |SRC| of Gain-Value Neurons in the gain context also became smaller as the wealth level increased. However, this trend did not reach a significant level (**Figure 3c**, right; permutation test; mean of |SRC_Gain_| in low wealth level = 2.38, mean of |SRC_gain_| in high wealth level= 2.06, p = 0.29; unsigned SRC in gain context in low or high wealth levels were indicated in light and dark green, respectively). Again, this wealth level-sensitive effect on AIC value coding (**Figure 3d**) is consistent with the fact that monkeys became less sensitive to objective value change when the wealth level increased (i.e., utility functions became flatter in both the loss and gain context when the wealth level increased; **Figure 1g**).

### Choice probability of AIC neurons predict the internal states related to behavioral choices

The gain/loss-specific value signals and the wealth level signals in AIC were present before the choice was made and were therefore in a position to influence the monkey’s decisions. To determine whether neural activity of the AIC neurons correlates with choice behavior, we computed a receiver operation characteristic (ROC) for each cell, and then computed the area under the curve (AUC) as a measure of the cell’s discrimination ability. In this analysis, an AUC value significantly different from 0.5 indicates at least a partial discrimination between two conditions. For each AIC neuron, we calculated two AUC values. First, we compared the firing rate distributions on choice trials when the monkey chose the gamble versus the sure option. We used this AUC value as a measure of *choice probability*. Second, we compared the firing rate distributions on choice trials when the monkey was risk-seeking versus risk-avoiding. We used this AUC value as a measure of *risk-attitude probability*. Risk-seeking trials were defined as trials where the monkey chose the gamble, even when the expected value of the gamble option was smaller than the expected value of the sure option. We did not include trials, in which the monkey chose the gamble option and it had a higher expected value, because in that case the monkey’s choice did not give any indication about his risk-attitude at that moment. Conversely, risk-avoiding trials were defined as trials where the monkey chose the sure option, even so it had a lower expected value than the gamble option. Thus, trials used to compute the risk-seeking probability were the subset of the trials used to compute choice probability, in which the monkeys did not make choices that maximized the expected value of the chosen option.

We found that trial-by-trial fluctuations in the activity of a subset of AIC neurons (35/240; 15%) significantly correlated with fluctuations of choice or risk-attitude. As shown in **Figure 4**, 19 neurons (8%) showed a significant choice probability (green), 20 neurons (8%) showed significant risk-attitude probability (purple), and 4 neurons (2%) showed both significant choice probability and risk-attitude probability (black). Across the AIC population (n=240), the ability of neural activity to predict choice and risk attitude showed a strong positive correlation (Pearson correlation; r = 0.41, p<10^−4^).

**Figure 4.**
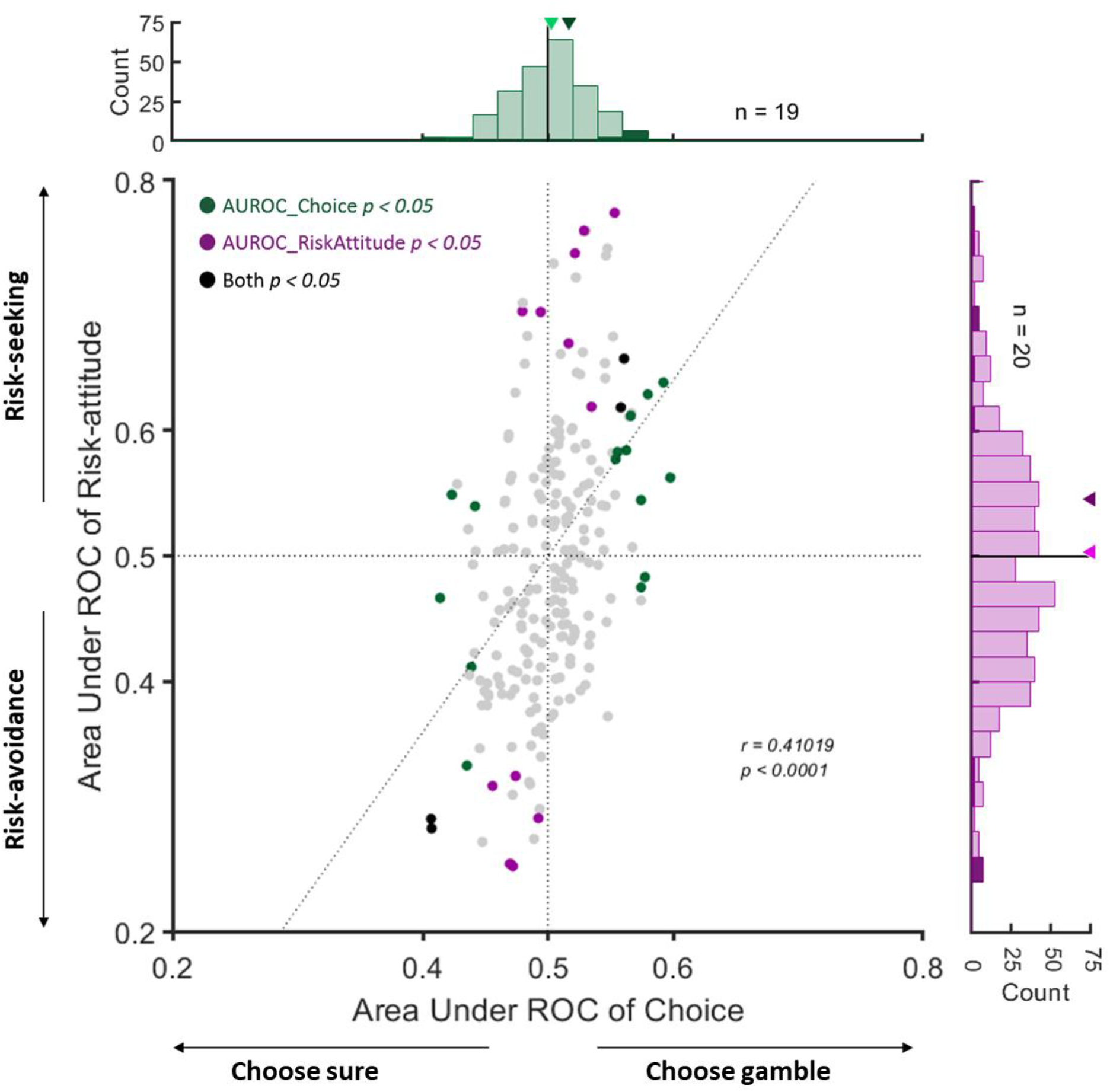
Distribution of area under the curve (AUC) of receiver operating characteristic (ROC) for choice and risk-attitude in individual neurons. AUC values capturing the covariation of each neuron with differences in choice (choosing gamble or sure) and risk-attitude (risk-seeking or risk-avoidance). Each point represents one neuron (n = 240), and colors indicate the significance of the two AUC values. In the marginal distributions, significant neurons are indicted in darker shades and the arrowheads indicate the average values across the entire distribution (light green or light purple) and the subset of neurons with significant AUC (dark green or dark purple), respectively. The gray vertical and horizontal dashed lines show the area of no significant discrimination ability (AUC of choice = 0.5 and AUC of risk-attitude = 0.5). The broken line represents the linear regression relating the AUC of choice and AUC of risk-attitude (*r* and *p* values refer to the regression slope).

AIC neurons encode contextual information that influence monkeys’ choice and momentary risk-attitude, such as current wealth level, gain/loss, and the value of each option (**Figure 2**). It is therefore not surprising that the activity of many of these neurons is predictive of choice or risk-taking. We examined whether neurons encoding specific-variables were particularly predictive of choice or risk-attitude (**Figure S6**). However, a chi-square test showed no significant dependency between the encoding of a specific decision-related variable and the likelihood that choice-predictive and/or risk-attitude-predictive signals were carried by a given AIC neuron (**Table3**, χ^2^= 7.61, p = 0.67, excluding neurons with AUROC that predict neither choice nor risk-attitude).

**Table 3.** The number of each signal during the choice period in the force choice trial, recounted based upon the AUC for choice or risk-attitude.

|  | Token + Value + Risk | Token + Value | Token + Risk | Value + Risk | Token | Value | Risk | End Token | None |
| --- | --- | --- | --- | --- | --- | --- | --- | --- | --- |
| Both | 0 | 0 | 1 | 0 | 3 | 0 | 0 | 0 | 0 |
| Choice | 0 | 0 | 3 | 0 | 5 | 0 | 2 | 1 | 4 |
| Risk-attitude | 0 | 1 | 3 | 0 | 8 | 0 | 3 | 0 | 1 |
| Neither | 3 | 12 | 23 | 1 | 104 | 5 | 16 | 9 | 32 |

## Discussion

Prospect theory provides profound insights into how humans make risky decisions in a wide range of circumstances^6,24^. The behavioral hallmarks described by the theory have also been reported in old- and new-world monkeys, as well as in rats^25–28^. This suggests that the neural circuits responsible for making risky decisions may have been evolutionarily conserved across mammals. Using a token-based gambling task, we demonstrated that activity of AIC neurons in macaques exhibits critical characteristics consistent with those postulated by Prospect Theory. Decoding the activity from a subgroup of AIC neurons can predict the monkey’s choices and risk attitude on a trial-by-trial basis. These results suggest that the AIC is a pivotal part of a circuit monitoring state and context information that controls risky choices by modifying activity in downstream decision processes.

Overall, monkeys in the present study were more prone to choose gamble options (**Figure. 1c**). This is in line with similar findings of previous monkey experiments^25,26,29,30^, yet is inconsistent to most human studies^6,24,31,32^. It is unclear whether such a discrepancy between humans and monkeys was due to species-specific differences, individual variability, or task design. Macaques have been shown to be risk-aversive, like humans, in a foraging task^33^ and in a risky decision-making task using animals’ hydration state to index their non-monetary wealth level^3^. The observed tendency to choose gamble options was therefore likely due to task-specific factors, such as the small reward amount at stake and the large number of trials.

Insular cortex is a large heterogeneous cortex that is typically divided into posterior granular, intermediate dysgranular, and anterior agranular sectors, based on cytoarchitectural differences. Our recordings were concentrated in the most anterior part of the insula and encompassed mostly agranular and some dysgranular areas (**Figure S4**). In addition, we also recorded some neurons in the border regions of the adjacent gustatory cortex. Importantly, we found no functional segregation or gradient with respect to the functional signals that were represented across the different cortical areas, we explored. This fits with a recent primate neuroimaging study that showed strong activation of this entire region by visual cues indicating reward, as well as reward delivery^20^. Insula has long been known to be strongly connected with the neighboring gustatory cortex^34,35^. Recently, several lines of studies have demonstrated that neurons in the gustatory cortex not only engage the primary processing of gustatory inputs, but also involve multisensory integration^15,16^, as well as higher cognitive functions like decision-making^13,35^. This suggests that primate insular cortex and the neighboring gustatory cortex are strongly interconnected and form an interacting distributed network during decision making.

Prospect theory assumes that people make decisions based on the potential gain or losses relative to a reference point. In our experiment, the natural reference point against which the monkey compared possible outcomes was the current token assets. Consistent with this idea, we found that the activity of a substantial number of AIC neurons (109/240; 45%) encoded start token number. The AIC has been suggested to represent the current physiological state of the subject (i.e., interoception)^15,16,36,37^. Our findings suggest that AIC also encodes more abstract state variables, such as current wealth level, which are important for economic decisions that will influence future homeostatic state. Notably, we found some AIC neurons encode the start token number in a numerical scale, with their activity increased (or decreased) specifically when the monkey owned a particular number of tokens. Such a pattern of numerical encoding has been identified in primate prefrontal and parietal cortex^38,39^, medial temporal lobe^40^, and recently in AIC^41^. It would be interesting for future studies to investigate whether these number-tuned neurons relate to the numerical abilities of primates.

Our results overwhelmingly support the notion that value-related signals in the brain operate in a relative framework. Only 4% of neurons in the AIC carried a value signal in an absolute framework. However, how the value of options is represented in a relative framework is an issue under debate. The core of the debate regards whether the value of gains and losses are represented in a single unitary system^42^ or separately by two independent systems^43–45^. Some of the AIC neurons encoding a parametric value signal did continuously across gains and losses (14/47; 30%). However, most AIC neurons encoded gain or loss-specific value signal (33/47; 70%). Thus, while there is some evidence for both hypotheses, most AIC value-encoding neurons form two independent representations that encode gains or losses, respectively. This functional separation is further supported by the presence of a large number of neurons carrying a categorical gain/loss signal. Interestingly, the number of loss-encoding neurons (29/33; 88%) is much larger than the number of gain-encoding AIC neurons (4/33; 12%). This may explain why human imaging studies often find a link between the AIC and the anticipation of aversive outcomes^1,13^. Thus, the AIC recordings show the presence of separate neuronal populations that encode value as a relative gain or loss. This could be the neuronal underpinning of the separate utility functions used in prospect theory.

‘Risk’ is often formalized and quantified as the outcome variance, and the AIC has been implicated to play a role in monitoring risk^22,23^. In line with this, we found 8% of the AIC neurons (n=19/240) whose activity correlates with the outcome variance. Moreover, the trial-by-trial variability of the monkeys’ choice and risk attitude was correlated with activity changes in a subgroup of AIC neurons (**Figure 6**). All of this supports the hypothesis that AIC plays an important part in the process of decision making under risk.

To the best of our knowledge, this is the first study recording single neurons of the AIC in awake, behaving primates during risky decision-making. We interpreted the function of AIC from the perspective of economic, risky decisions ^13^ and within the framework of Prospect theory^6^. Decisions are likely not only guided by the rational, abstract processes depicted by economic models, but are strongly influenced by emotional processes^46^. The AIC has been suggested to occupy a central position in regulating emotions as it receives interoceptive afferents from visceral organs through the posterior granular insula area, representing contextual information (as demonstrated by this study), and is closely connected with the amygdala and autonomic nuclei^15,47^. This study took the first step to delineate how the decision context modulates economic value representation, and thereby impacts the decision of subjects. Future work will further investigate the interacting functions of AIC in economic decisions, emotions, and autonomic regulation.

## Materials and Methods

All animal care and experimental procedures were conducted in accordance with the US public Health Service policy on the humane care and use of laboratory animals and were approved by the Johns Hopkins University Institutional Animal Care and Use Committee (IACUC).

### General

Two male rhesus monkeys (Monkey G: 7.2 kg, Monkey O: 9.5 kg) were trained to perform a token-based gambling task in this study. Monkeylogic software^48^ (https://www.brown.edu/Research/monkeylogic/) was used to control task events, stimuli, and reward, as well as monitor and store behavioral events. During the experimental sessions, the monkey was seated in an electrically insulated enclosure with its head restrained, facing a video monitor. Eye positions were monitored with an infrared corneal reflection system, EyeLink 1000 (SR Research) at a sampling rate of 1000 Hz. All analyses were performed using self-written Matlab code, unless noted otherwise.

### Behavioral tasks

The token-based gambling task was based on a previously published task design^49^ and consisted of two types of trials: choice trails and force choice trials. In choice trials, two targets (both a sure option and a gamble option) were presented on the screen. Monkeys were allowed to choose one of the options by making a saccade to the corresponding target. Choice trials allowed us to measure the monkey’s risk attitude in different behavioral contexts of various value-matching of gamble and sure. In force choice trials, only one target (either a sure option or a gamble option) was presented on the screen so the monkey was forced to make a saccade to the given target. Comparing the neuronal activity in choice and force choice trials allowed us to identify neuronal signals specifically related to decision-making. The choice and force choice trials were pseudo-randomly interleaved in blocks so that each block consisted of all 24 choice trials and 13 force choice trials.

A choice trial began with the appearance of a fixation point surrounded by the token cue. After the monkey had maintained its gaze at the central fixation point (±1° of visual angle) for a delay period (0.5-1s), two choice targets were displayed on two randomly chosen locations among the four quadrants on the screen. The monkey indicated its choice by shifting gaze to the target. Following the saccade, the token cue moved to surround the chosen target and the unchosen target disappeared from the screen. The monkey was required to keep fixating the chosen target for 450-550ms, after which the chosen target changed either color or shape. If the chosen target was a gamble option, it changed from a two-colored square to a single-colored square to indicate the outcome of the gamble. The color represented the amount of gained or lost tokens in the present trial. If the chosen target was a sure option, the shape changed from a square to a circle serves as a control to the change in visual display that occurs during gamble option choices. Finally, after an additional delay (500ms) the token cue was updated. If the owned token number was equal to or more than 6 at this stage, the monkey received a standard fluid reward after an additional 450ms waiting time. At the beginning of the next trial, the remaining tokens were displayed with filled circles. Otherwise, if the owned token number was smaller than 6, the monkey did not receive a fluid reward and the updated token cue was displayed at the beginning of the next trial. If the owned token number was smaller than 0, the inter-trial-interval (ITI) for the next trial would be prolonged (300 ms per owed token).

The monkey was required to maintain the fixation spot until it disappeared for reward delivery. If the monkey broke fixation in either one of the two time periods, the trial was aborted and no reward was delivered. The following trial repeated the condition of the aborted trial contingent on the time of fixation break. A trial in which the monkey broke fixation *before* the choice was followed by a trial in which the same choice targets were presented, but at different locations. This ensured that the monkey sampled every reward contingency evenly but could not prepare a saccade in advance. On the other hand, a trial in which the monkey broke fixation *after* the choice was followed by a no-choice trial in which only the chosen target was presented. If the monkey broke fixation following a gamble choice, but before the gamble outcome was revealed, the same gamble cue was presented. If the monkey broke fixation following a sure choice or after a gamble outcome was revealed, the same sure cue was presented. This ensured that the monkey could not escape a choice once it was made and had to experience its outcome. All trials were followed by a regular 1500-2000ms ITI. The schedule of the token-based gambling task is shown in **Figure 1a**.

All options in this task were represented by sets of colored squares, with the color of the square indicating the token amount that could be gained or lost (token outcome) and the proportion of color indicating the probability that this event would take place (outcome probability) (**Figure 1b**). The sure options were single-colored squares indicating a certain outcome (gain or loss of token). There were 7 different colors used for sure options representing the number of tokens that were gained or lost ([−3, −2, −1, 0, +1, +2, +3]). The gamble options were two-colored squares indicating two possible outcomes indicated by two different colors. The portion of each color corresponded to the probability of each outcome. Six gamble options were used in this task. Three of the gambles resulted in either a gain of 3 or 0 token(s), but with different outcome probabilities (i.e. token [+3, 0] with the probability combination of [0.1, 0.9], [0.5, 0.5], or [0.75, 0.25]). Another three gambles resulted either in a loss of 0 or 3 token(s) with different outcome probability (i.e. token [0, −3] with probability combination of [0.1, 0.9], [0.5, 0.5], or [0.75, 0.25]). The choice trials were divided into a gain context and a loss context (**Figure 1b**). In the gain context, the three gamble options that resulted in either a gain of +3 or 0 token with different outcome probabilities were paired with four sure options that spanned the range of gaining outcomes (i.e. [0, +1, +2, +3]). These resulted in 12 possible combinations of sure and gamble options. In the loss context, the other three gamble options resulted either in a loss of 3 or 0 with probability combination were paired with four sure options that spanned the range of losing outcomes (i.e. [0, −1, −2, −3)). Thus, there were another 12 possible combinations of sure and gamble options. This resulted in a total of 24 different combinations of reward option combinations (half in the gain context and the other half in the loss context) that are offered in choice trials. In the force choice trials, all 13 different reward options (7 sure and 6 gamble options) which were used in the choice trials are presented in isolation.

### Saccade detection

Eye movements were detected offline using a computer algorithm (saccade detection function) that searched first for significantly elevated velocity (30°/s). Saccade initiations were defined as the beginning of the monotonic change in eye position lasting 15ms before the high-velocity gaze shift. A valid saccade for choice was further admitted to the behavioral analysis if it started from the central fixation window (1° × 1° of visual angle) and ended in the peripheral target window (2.5° × 2.5° of visual angle).

### Description of monkeys’ behavior

Fixation behavior: We examined whether and how monkeys’ motivations to initiate a new trial were influenced by the outcome of the previous trial and the start token number of the current trial. We used two behavioral signals as indications of the monkey’s motivational state: (1) fixation latency (i.e., the time from fixation point onset until fixation by the monkey) and (2) fixation break ratio (i.e., the frequency with which the monkey failed to fixate on the fixation point long enough to initiate target onset). We used linear regression models to test if there was a significant relationship between each of the two variables describing motivational state and the variables describing history and current state.

Response time: We examined whether and how response times were influenced by different decision-related variables. For each trial, response time was defined as the time period between target onset and saccade initiation estimated by the saccade detection function. The response time dataset in each condition (e.g. trials from context of gain or loss, trials with different start token numbers, trials with different expected values of chosen option (chosen EV), or trials with different absolute values of difference of expected values among the gamble and sure option (|ΔEVgs|)) was fitted with an ex-Gaussian distribution algorithm^50^ (https://doi.org/10.6084/m9.figshare.971318.v2). It returned three besting-fitting parameter values of the ex-Gaussian distribution: the mean μ, the variance σ, and the skewness τ of the distribution. We used a permutation test to determine if the mean RTs of trials from the gain and loss context. We used linear regression models to test whether there was a significant relationship between mean RTs and start token number, chosen EV, or |ΔEVgs|.

### Prospect theory model

Prospect theory is derived from classical expected value theory in economics^51^ and assumes that the subjective value of a gamble depends on the utility of the reward amount that can be earned, weighted by the ‘subjective’ estimation of the probability of the particular outcome. Both the utility function and the probability function can be non-linear and thus might influence risk preference.

We modeled the probability that monkeys chose the gamble option by a softmax choice function whose argument was the difference between the subjective values of each option.

Subjective utility was parameterized as:

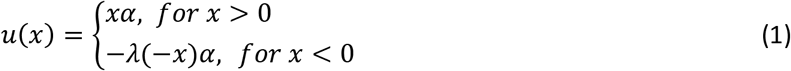

where α is a free parameter determining the curvature of the utility function, u(x), and x is the reward outcome (in units of gaining or losing token numbers).

Subjective probability of each option is computed by:

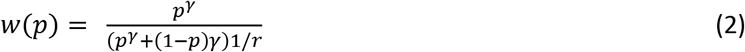

where γ is a free parameter determining the curvature of the probability weighting function, w(p), and p is the objective probability of receiving corresponding outcome. The u(x) and w(p) were followed with research^2,6^.

The subjective value (SV, or say expected utility) of each option was computed by combining the output of u(x) and w(p) that map objective gains and losses relative to the reference point and objective probability onto subjective quantities, respectively:

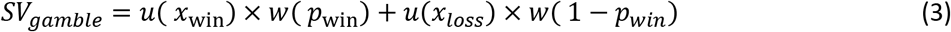

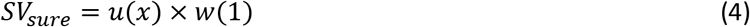

The subjective value difference between the two options was then transformed into choice probabilities via a softmax function with terms of slope s and bias s:

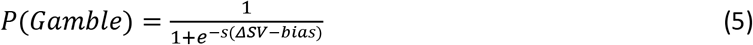

where *ΔSV* = *SV*_*gamble*_ − *SV*_*sure*_, *s*, determines the sensitivity of choices to the ΔSV, and is the directional bias of choosing gamble.

For an alternative expected value (EV) model, the value of option is calculated as:

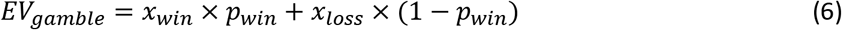

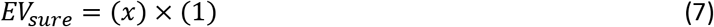

The expected value difference between the two options was then transformed into choice probabilities via a softmax function with terms of slope s and bias s as what we did in the PT model:

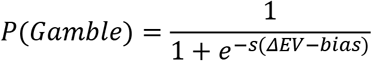

where *ΔEV* = *EV*_*gamble*_ − *EV*_*sure*_, *s*, determines the sensitivity of choices to the ΔSV, and is the directional bias of choosing gamble.

We optimized model parameters, *α*, *λ*, *γ*, in the PT model, and *s* and *bias* and in both PT and EV models by minimize the negative log likelihoods of the data given different parameters setting using Matlab’s fmincon function, initialized at multiple starting points of the parameter space as follows:

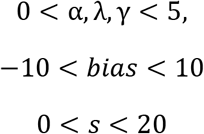

−2 negative log-likelihoods (−2\**LL*_*max*_ which measures the accuracy of the fit) were used to compute classical model selection criteria. We also computed the Bayesian information criterion (BIC):

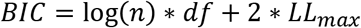

where *n* is the number of training trial and *df* is the number of free parameters in the model. The likelihood in BIC is penalized by adding more parameters into the model. Thus, we use BIC to represent the trade-off between model accuracy and model complexity and use it to guide model selection. We then compared −2\**LL*_*max*_ and BIC calculated from a 5-fold cross-validation with separate training and testing for the PT model and EV model in paired *t-tests*. We also generated model simulations for PT and EV model in **Figure S2** after optimizing model’s parameters.

As in classical expected value theory in economics^52^, a convex utility function (α > 1) implies risk seeking, because in this scenario, the subject values large reward amounts disproportionally more than small reward amounts. Gain from winning the gamble thus has a stronger influence on choice than loss from losing the gamble. In the same way, a concave utility function (α < 1) implies risk seeking, because large reward amounts are valued disproportionally less than small ones.

The *λ* can further influence subject’s risk-attitude in the context of gain or loss because it modulate the curvature of utility function in gain to that in the loss context. With a *λ* > 1, the utility function in the loss context will be sharper than that in the gain context, indicating the subject is more sensitive to the value change in the loss context. While with a *λ* < 1, the utility function in the loss context will be flatter than that in the gain context, indicating the subject is less sensitive to the value change in the loss context.

Independently, a non-linear weighting of probabilities can also influence risk attitude. For example, an S-shaped probability weighting function (*γ* < 1) implies that the subject overweighs small probabilities and underweights the large probabilities. This would lead to higher willingness to accept a risky gamble, because small probabilities to win large amounts would be overweighed relative to high probabilities to win moderate amounts.

The bias term in the softmax function can also influence a subject’s risky choices independent to the subjective value of options. A negative bias will result with risk-seeking behavior because the subject tends to choose gamble while the SVs of gamble and sure are identical. In the other hand, a positive bias will result with risk-aversive behavior because the subject tends to choose sure while the SVs of gamble and sure are identical.

### Cortical localization and estimation of recording locations

We used T1 and T2 magnetic resonance images (MRIs) obtained for the monkey (3.0 T) to determine the location of the anterior insula. In primates, the insular cortex constitutes a separate cortical lobe, located on the lateral aspect of the forebrain, in the depth of the Sylvian or lateral fissure (LF) (**Figure 2g**). It is adjoined anteriorly by the orbital prefrontal cortex, and it is covered dorsally by the frontoparietal operculum and ventrally by the temporal operculum. The excision of the two opercula and part of the orbital prefrontal cortex reveals the insula proper, delimited by the anterior, superior, and inferior peri-insular (or limiting or circular) sulci. We used the known stereoscopic recording chamber location and recording depth of the electrode to estimate the location of each recorded neurons. The estimated recording locations were superimposed on the MRI scans of each monkey. Cortical areas were estimated using the second updated version of the macaque monkey brain atlas by Saleem and Logothetis^54^ with a web-based brain atlases^55^.

### Surgical procedures

Each animal was surgically implanted with a titanium head post and a hexagonal titanium recording chamber (29mm in diameter) 20.5 mm (Monkey G) and 16 mm (Monkey O) lateral to the midline, and 30 mm (Monkey G) and 34.5 mm (Monkey O) anterior of the interaural line. A craniotomy was then performed in the chambers on each animal, allowing access to the AIC. The location of AIC was determined with T1 and T2 magnetic resonance images (MRIs, 3.0T) obtained for each monkey. All sterile surgeries were performed under anesthesia. Post-surgical pain was controlled with an opiate analgesic (buprenex; 0.01 mg/kg IM), administered twice daily for 5 days postoperatively.

### Neurophysiological recording procedures

Single neuron activities were recorded extracellularly with single tungsten microelectrodes (impedance of 2-4 MΩs, Frederick Haer, Bowdoinham, ME). Electrodes were inserted through a guide tube positioned just above the surface of the dura mater and were lowered into the cortex under control of a self-built Microdrive system. The electrodes penetrated the cortex perpendicular to the surface of the cortex. The depths of the neurons were estimated by their recording locations relative to the surface of the cortex. Electrophysiological data were collected using the TDT system (Tucker & Davis). Action potentials were amplified, filtered, and discriminated conventionally with a time-amplitude window discriminator. Spikes were isolated online if the amplitude of the action potential was sufficiently above a background threshold to reliably trigger a time-amplitude window discriminator and the waveform of the action potential was invariant and sustained throughout the experimental recording. Spikes were then identified using principal component analysis (PCA) and the time stamps were collected at a sampling rate of 1,000 Hz.

### Spike density function

To represent neural activity as a continuous function, we calculated spike density functions by convolving the peri-stimulus time histogram with a growth-decay exponential function that resembled a postsynaptic potential^56^. Each spike therefore exerts influence only forward in time. The equation describes rate (R) as a function of time (t): R(t) = (1 - exp(-t/τg))*exp(-t/τd), where τg is the time constant for the growth phase of the potential and τd is the time constant for the decay phase. Based on physiological data from excitatory synapses, we used 1 ms for the value of τg and 20 ms for the value of τd^57^.

### Linear regression analysis of neuronal coding

To find AIC neurons, whose neuronal activities reflect specific decision-related variable(s), we performed a linear regression with its mean Firing rate (FR) within the choice period for each trial as the dependent variable, and a predictor derived from the decision-related variables as the independent variable:

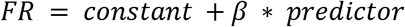

in which, β was the coefficient of the predictor. A constant term was added as a baseline model.

To tested four different classes of decision-related variables: (1) “Token-asset” variables, (2) “Gain/Loss-Value” variables, (3) “Risk” variables, and (4) a “Absolute value” variable. In total, we considered 16 different potential decision-related variables, as well as a baseline model that consisted only of a constant term.

Token-asset variables were variables that represented the start token number in one of three different types. The first type, the *linear token signal*, encoded the start token number in a linear, continuous manner (monotonically rising or falling from 0 to 5). The second type, the *binary token signal*, encoded the start token number in a binary, discontinuous manner (with a value of “1” for trials with start token number 0 to 3 and a value of “2” for trials with start token number 3-5). The third type, the *numerical token signal*, encoded the start token number in a Gaussian manner with the peak of the activity at one of the token numbers from 0 to 5, and the activity symmetrically falling for token numbers that were smaller or larger than the peak.

Gain/Loss-Value variables were variables that represented the gain/loss context, the expected value of options, or gain/loss context-dependent value signals. We tested five types of variables. The first type, the *gain/loss signal*, encoded the context of gain or loss in a binary manner. Trials in the gain context were indicated with a “1”, and trials in the loss context with a “-1”. The only exception were no-choice trials with a sure option with EV = 0, which were indicated with a “0”. The second type, the *linear value signal*, encoded the expected value of options in a linear, continuous manner across both the gain and loss context (with a range from −3 to 3). The remaining types also encoded the expected value of options, but contingent on the gain/loss context. The third type, the *gain value signal*, encoded the expected value of options in a linear manner, but only in the gain context (options with EV larger than 0 were encoded as the original number, otherwise were encoded as “0”), while the fourth type, the *loss value signal*, encoded the expected value of options in a linear manner only in the loss context (options with EV smaller than 0 were encoded as the original number, otherwise were encoded as “0”). The fifth type, the *behavioral salience signal*, encoded the expected value of options in a linear but asymmetric direction for the gain and loss context. Thus, this signal encoded the absolute distance of the value from zero, independent of whether it represented a gain or a loss (e.g. both an option with EV = 1.5 and an option with EV = −1.5 would be encoded as “1.5”).

Risk variables were variables that represented the variance of possible outcomes of an option (calculated by 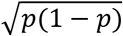, in which p was the winning probability of the option). We considered two different types. The first type, the *linear risk signal*, encoded the variance of outcome in a linear manner proportional to the variance. The second type, the *binary risk signal*, encoded whether the outcome of option was uncertain or not in a binary manner (with a value of “1” for all gamble options and a value of “0” for all sure options).

So far, value always was defined as a relative change of tokens, independent of the start token number. We also considered an a*bsolute value signal* (i.e., the sum of all possible end token numbers, weight by their probability). This signal took into account not only the possible change in token number, but also the start token number. Thus, it represented the outcome of a choice in absolute framework that reflected how close the monkey was to earning fluid reward.

### Mix-selective neuronal coding with regression analysis

The regression analysis using the series of single variable models indicated that many neurons encoded multiple decision-related variables. We therefore further investigated the contribution of decision-related variables to neural activity, by using a multiple linear regression with the mean Firing rate (FR) within the choice period for each trial as the dependent variable, and predictors derived from the decision-related variables as the independent variables.

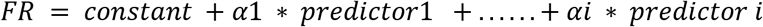

We fitted a family of regression models with all possible combination of the basic decision-related variables described in the last section. This resulted in a total of 163 tested models. For each neuron, we determined the best-fitting model using the Akaike information criterion and classified it as belonging to different functional categories according to the variables that were included in the best-fitting model.

### Sensitivity to value changes in gain and loss value cells

We examined whether neurons carrying *Loss value signals* and *Gain Value signals* showed different sensitivity to changes in value. We estimated the absolute value of the standardized regression coefficients (|SRCs|) of firing rate of Loss-Value Neurons in the loss context and |SRCs| of firing rate of Gain-Value Neuron in the gain context, respectively. We included all neurons for statistical test, whose best-fitting model included a Loss or Gain Value Signal. We performed a permutation test with 10,000 iterations to test if the normalized |SRCs| of Loss Value (n = 39) and Gain Value neurons (n = 12) showed a significant difference (**Figure 3a**).

We also examined whether the sensitivity to value change in neurons carrying *Loss value signals* and *Gain Value signals* was influenced by the wealth level (i.e., the number of tokens owned at the beginning of the trial). We compared |SRCs| of Loss and Gain Value neurons in trials with low [0,1,2] or high [3,4,5] token level. We performed a permutation test with 10,000 iterations to test if the normalized |SRCs| of Loss Value (n = 39) and Gain Value neurons (n = 12) showed a significant difference for low and high token levels (**Figure 3c**).

### Receiver operating criterion (ROC) analysis

To determine whether neural activity of the AIC neurons was correlated with the monkey’s choice behavior or risk-attitude, we computed a receiver operation characteristic (ROC) for each cell and computed the area under the curve (AUC) as a measure of the cell’s discrimination ability. We computed the AUC value of *choice probability* by comparing the distributions of firing rates associated with each of the two choices (i.e. choice of “gamble” or choice of “sure”). We computed the AUC value of *risk-seeking probability* by comparing the two distributions of firing rates associated with risk-seeking and risk-avoidance behavior. Risk-seeking trials were defined as trials where the monkey chose the gamble, even so the expected value of the gamble option was smaller than the expected value of the sure option. We did not include trials, in which the monkey chose the gamble option and it had a higher expected value, because in that case the monkey’s choice did not give any indication about his risk-attitude at that moment. Conversely, risk-avoiding trials were defined as trials where the monkey chose the sure option, even so the expected value of the sure option was smaller than the expected value of the gamble option. Thus, trials used to compute the risk-seeking probability were a subset of the trials used to compute choice probability, in which the monkeys did not make choices that maximized the expected value of the chosen option.

### Chi-square test

To test whether neurons encoding specific behaviorally relevant variables were more likely to carry significant choice or risk-attitude probability signals, we used a chi-square test, which is used to determine whether there is a statistically significant difference between the expected frequencies and the observed frequencies in one or more categories.

## Supplementary materials

**Figure S1.**
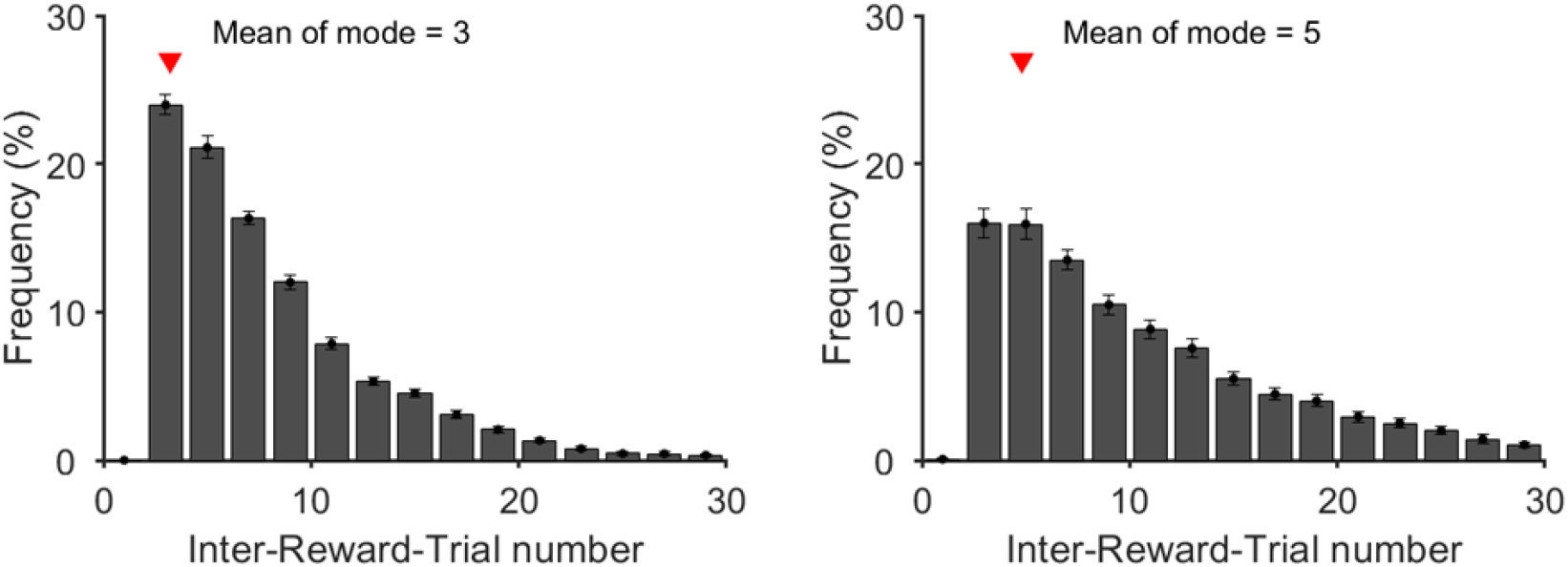
Inter-Reward-Trial number for each monkey. Monkeys used to accumulate the necessary 6 tokens for a standard fluid reward in 3-5 successive trials. Red triangle indicates the average trials to get the reward (I think it is better to use a vertical line to indicate average trials). Error bars indicate SEM or estimates across sessions (session number = 37 for each monkey).

**Figure S2.**
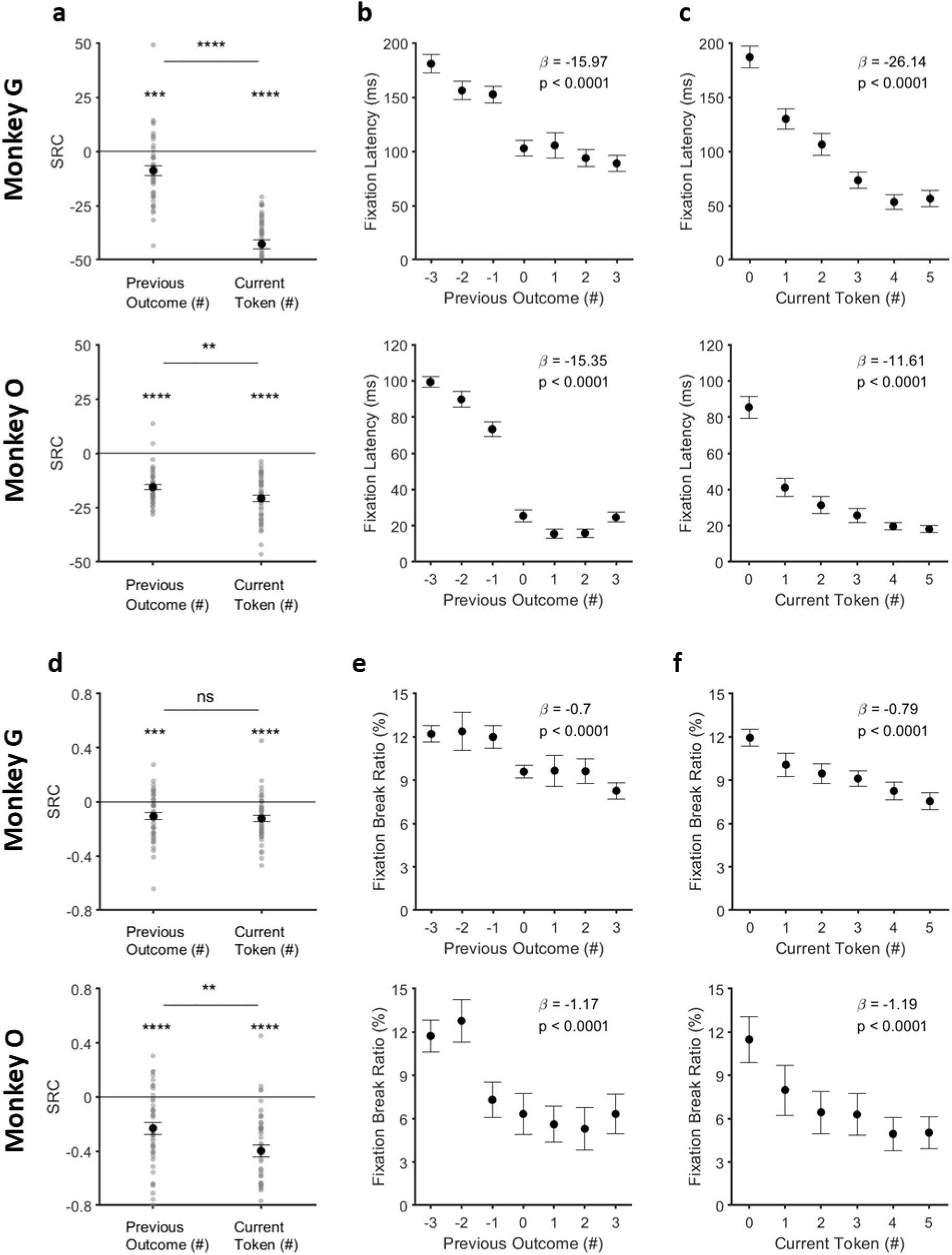
Effect of current token asset and token outcome history on fixation latency and fixation break ratio. (a) Standardized regression coefficients (SRCs) for fixation latency (latency to fixate on the center point at the beginning of the trial before the token cue appears) as a function of previous outcome (token change in last trial) and current token number. Error bars indicate SEM of SRCs across sessions. (b) Fixation latency as a function of previous outcome. Regression coefficient (β) between fixation latency and the token number won or losses of the previous trial. (c) Fixation latency as a function of current token asset. Regression coefficient (β) between fixation latency and the start token number of the current trial. (d) Standardized regression coefficients (SRCs) for fixation break ratio (failure to hold fixation on the center point long enough for token cues to appear) as a function of previous outcome (token change in last trial) and current token number. Error bars indicate SEM of SRCs across sessions. (e) Fixation break ratio as a function of previous outcome. Regression coefficient (β) between percentage of trials with fixation breaks and the token number won or losses of the previous trial. (f) Fixation break ratio as a function of current token asset. Regression coefficient (β) between percentage of trials with fixation breaks and the start token number of the current trial. Error bars indicate SEM or estimates across sessions (session number = 37 for each monkey). ns, no significant, ** p<10^−2^, *** p<10^−3^, **** p<10^−4^ (t-test or paired t-test).

**Figure S3.**
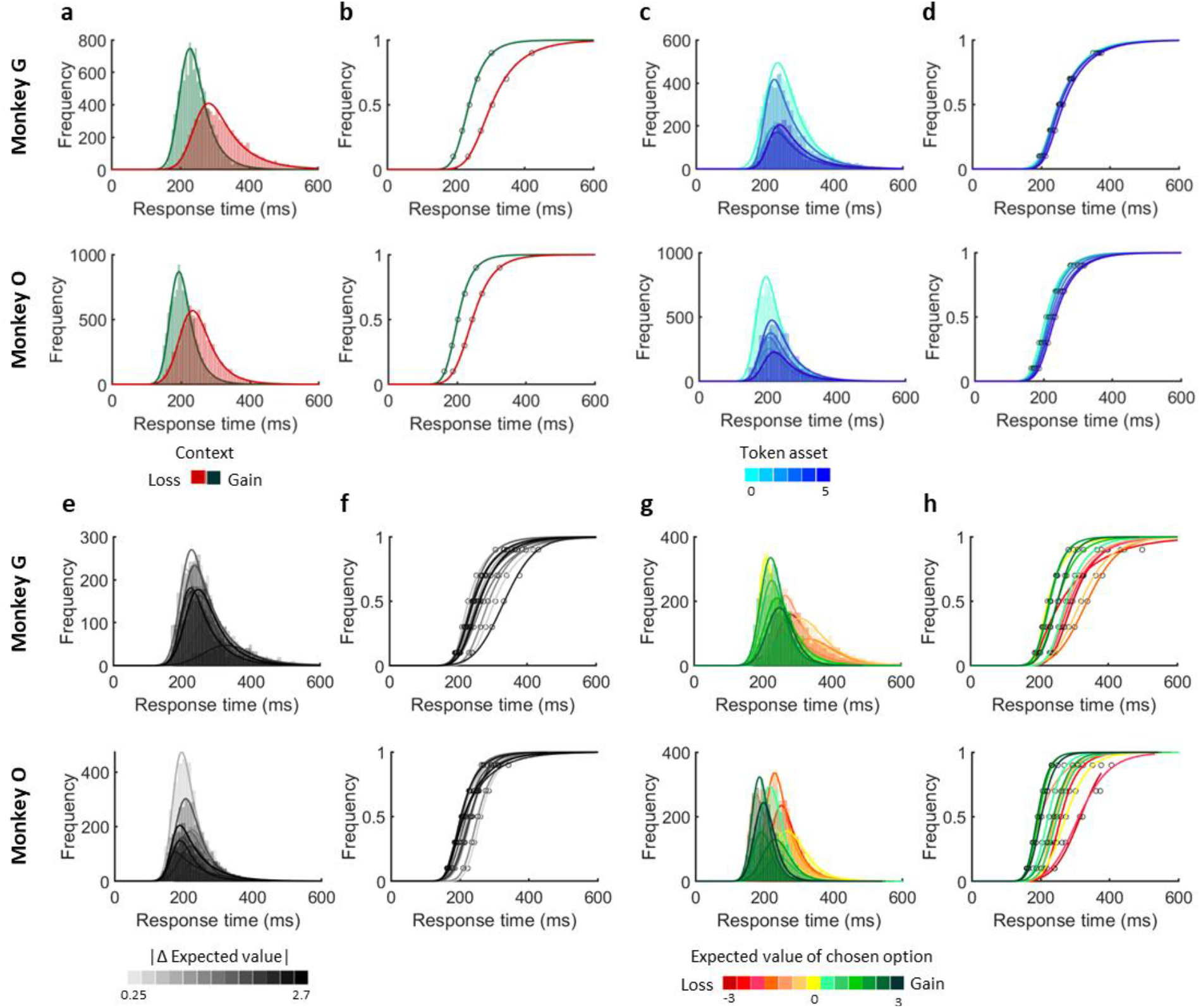
Response time of monkey’s choice. (a) Distribution of response times when monkeys made decisions in the gain (green) and loss (red) context for each monkey. Histograms with light color indicate the raw data distribution and curves with dark color indicate the best-fitting (ex-Gaussian) distribution. (b) Cumulative distribution function (CDF) of response times in the gain (green) and loss (red) context. (c) Distribution of response times when monkeys made decisions with different start token numbers. The color gradients from light to dark blue indicate token number from 0 to 5. (d) CDF of response times with different start token numbers. (e) Distribution of response times when monkeys made decisions with different absolute values of expected value difference between gamble and sure option (|ΔEVgs|s). The black color gradients indicate |ΔEVgs| from small to large. (f) CDF of response times with different |ΔEVgs|s). (g) Distribution of response times when monkeys made decisions with different EV of chosen option. The color gradients indicate chosen values from small to large. (h) CDF of response times with different EV of chosen option. One monkey took more time to make a choice when the difference expected value between options were small (Figure S3e-f; regression analysis; monkey O: β _RT_|ΔEVgs|_= −5.11, p < 10^−3^), indicating a task-difficulty dependent response time. Yet another monkey showed no significant difference to this variable (Figure S3e-f; regression analysis; monkey G: β _RT_|ΔEVgs|_= −0.50, p = 0.74). Furthermore, Both monkeys made faster choices as the expected value of chosen option increased (Figure S3g-h; regression analysis; monkey G: β _RT_StartTkn_= 2.83, p = 0.19; monkey O: β _RT_StartTkn_= 3.50, p < 10^−2^). This likely reflects an elevated motivation for high-value options.

**Figure S4.**
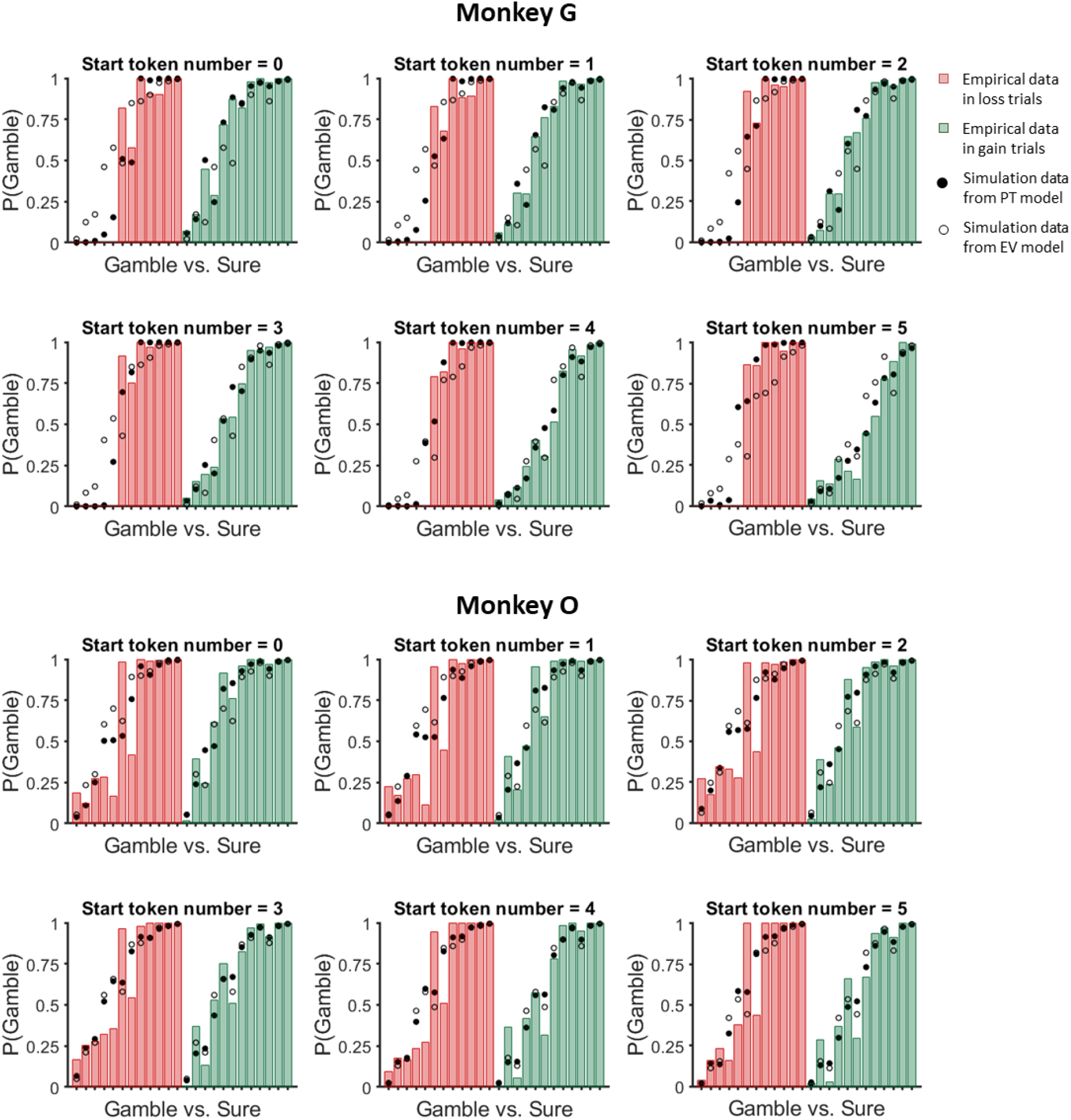
Behavioral results and model simulations. Probability of choosing gamble, P(Gamble) for all possible choice target pairs (a gamble and a sure from gain or loss context) correspond to token asset 0-5. Colored bars represent the actual choice data and black (prospect theory model) and white (expected value model) dots represent the model simulated data. Bars are sorted according to ΔEV (EV gamble - EV sure) from small to large in loss and gain context, respectively.

**Figure S5.**
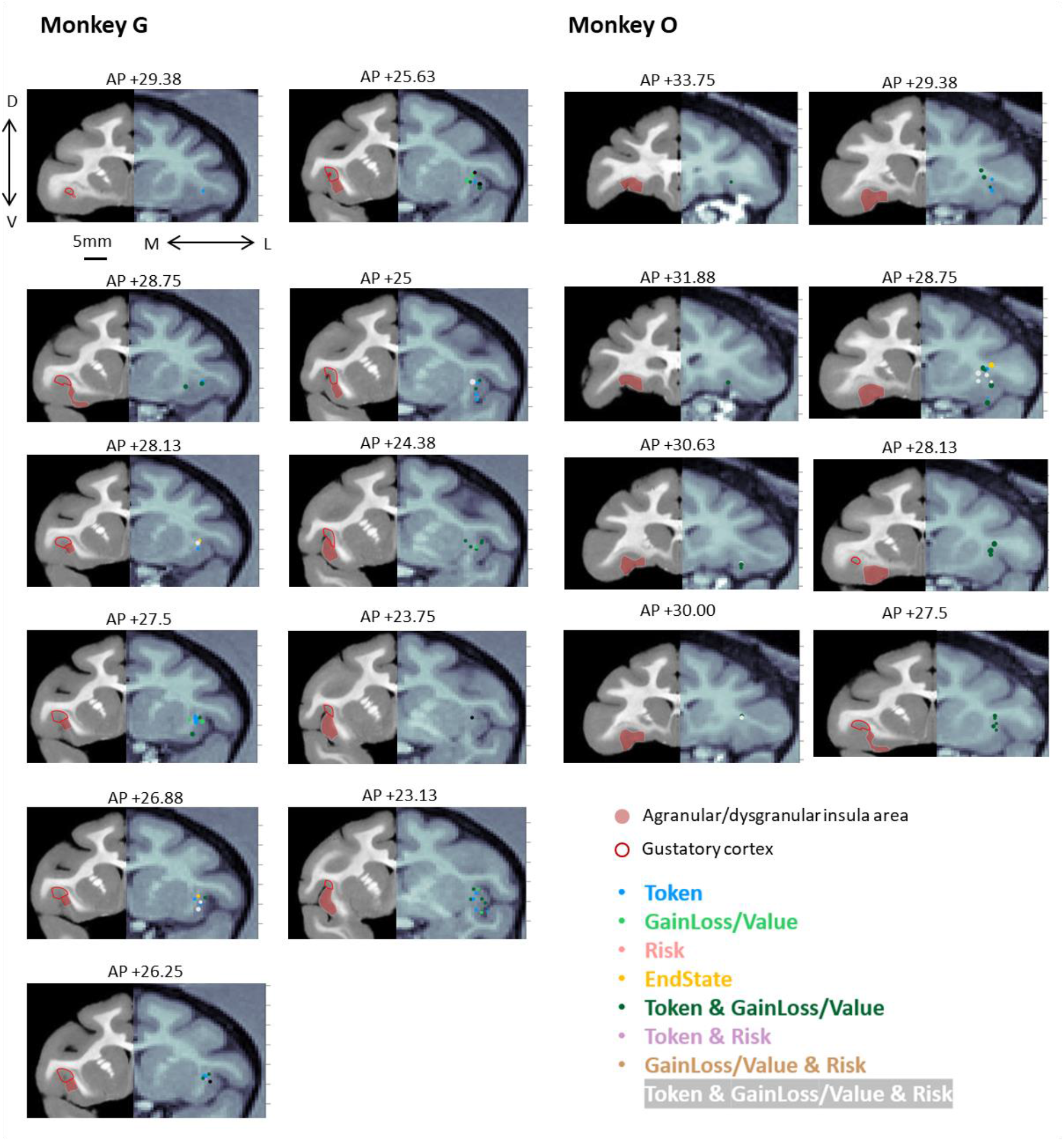
Recording sites with location of neurons of different functional types. Coronal MRI sections for each monkey show the locations of recorded neurons. The right side of each section shows the MRI from the anatomical scan of each monkey performed before surgery. Superimposed on each section is the estimated location of each recorded neuron based on penetration coordinates and recording depth. Neuronal classification according to the regression model is marked in different colors. The dot size indicates the number of units recorded in the location. Different colors indicate different functional signals encoded by the neurons. The position of each section in stereotactic coordinates is indicated on top. The left side of each section shows the most similar section in the macaque brain atlas of Saleem and Logothetis (2012). The location of the agranular and dysgranular insula (filled pink area), and gustatory cortex (red outlined area) are indicated in each section.

**Figure S6.**
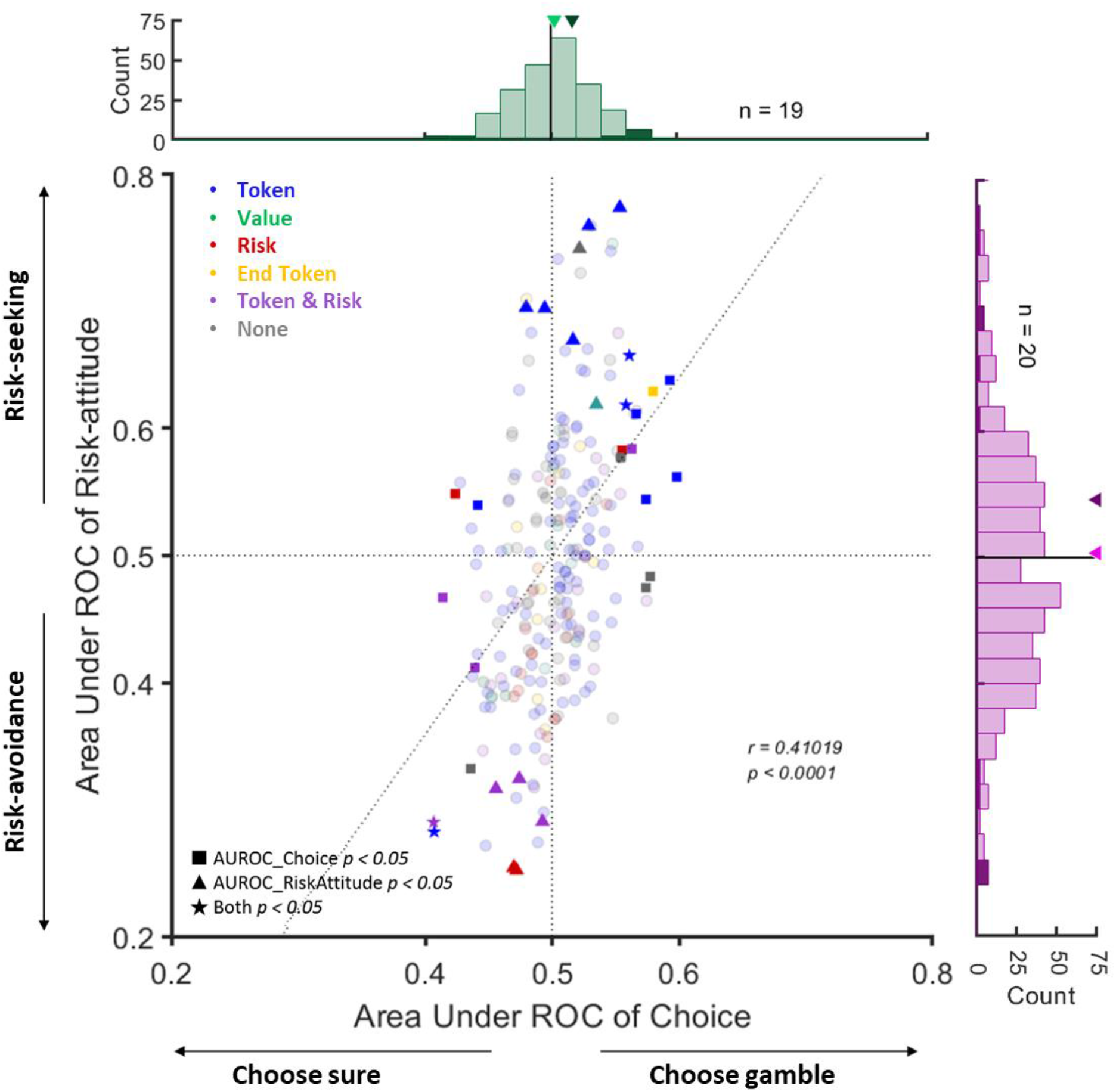
Distribution of area under the curve (AUC) of receiver operating characteristic (ROC) for choice and risk-attitude in neurons encode different kind of decision-related signals. AUC values capturing the covariation of each neuron with differences in choice (choosing gamble or sure) and risk-attitude (risk-seeking or risk-avoidance). Each point represents one neuron (n = 240). Shapes indicate the significance of the two AUC values. Colors indicate the functional signal encoded by the neuron. In the marginal distributions, significant neurons are indicted in darker shades and the arrowheads indicate the average values across the entire distribution (light green or light purple) and the subset of neurons with significant AUC (dark green or dark purple), respectively. The gray vertical and horizontal dashed lines show the area of no significant discrimination ability (AUC of choice = 0.5 and AUC of risk-attitude = 0.5). The broken line represents the linear regression relating the AUC of choice and AUC of risk-attitude (*r* and *p* values refer to the regression slope).

**Table S1.**
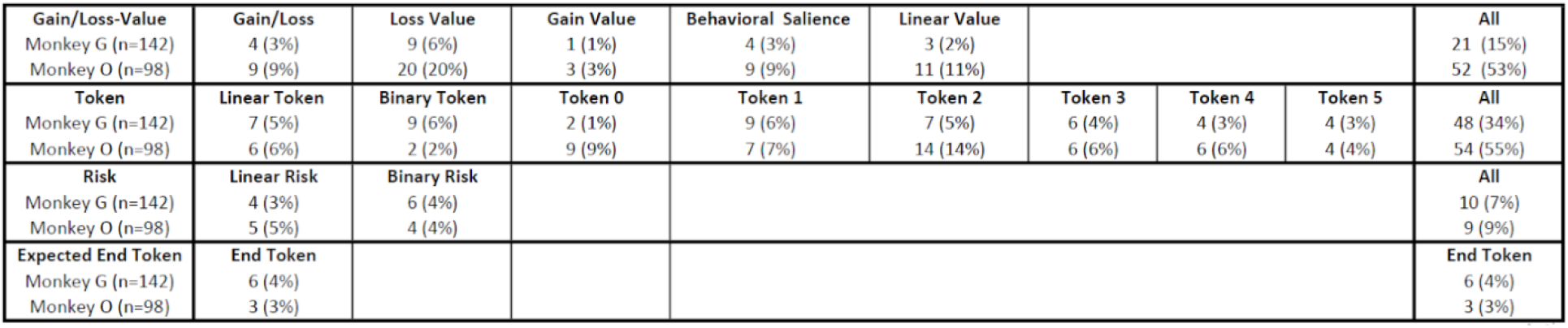
Summary of the number and percentage of significant responding neurons in different subsets of neuron types for AIC neurons recorded from each monkey.

**Table S2.** Summary of the number and percentage of neurons positively or negatively correlated to different decision-related variables.

| Gain/Loss-Value | Gain/Loss | Loss Value | Gain Value | Behavioral Salience | Linear Value |  |  |  | All |
| --- | --- | --- | --- | --- | --- | --- | --- | --- | --- |
| Positive correlation | 8 (62%) | 27 (93%) | 0 (0%) | 0 (0%) | 13 (93%) |  |  |  | 48 (66%) |
| Negative correlation | 5 (38%) | 2 (7%) | 4 (100%) | 13 (100%) | 1 (7%) |  |  |  | 25 (34%) |
| Token | Linear Token | Binary Token | Token 0 | Token 1 | Token 2 | Token 3 | Token 4 | Token 5 | All |
| Positive correlation | 11 (85%) | 4 (36%) | 0 (0%) | 8 (50%) | 21 (88%) | 12 (100%) | 8 (80%) | 5 (63%) | 69 (66%) |
| Negative correlation | 2 (15%) | 7 (64%) | 11 (100%) | 8 (50%) | 3 (12%) | 0 (0%) | 2 (20%) | 3 (37%) | 36 (34%) |
| Risk | Linear Risk | Binary Risk |  |  |  |  |  |  | All |
| Positive correlation | 3 (33%) | 3 (30%) |  |  |  |  |  |  | 6 (32%) |
| Negative correlation | 6 (64%) | 7 (70%) |  |  |  |  |  |  | 13 (68%) |
| Expected End Token | End Token |  |  |  |  |  |  |  | End Token |
| Positive correlation | 5 (56%) |  |  |  |  |  |  |  | 5 (56%) |
| Negative correlation | 4 (44%) |  |  |  |  |  |  |  | 4 (44%) |

## Reference

1. Canessa, N. et al. The functional and structural neural basis of individual differences in loss aversion. J. Neurosci. 33, 14307–14317 (2013).

2. Juechems, K., Balaguer, J., Ruz, M. & Summerfield, C. Ventromedial prefrontal cortex encodes a latent estimate of cumulative reward. Neuron 93, 705–714. e4 (2017).

3. Yamada, H., Tymula, A., Louie, K. & Glimcher, P. W. Thirst-dependent risk preferences in monkeys identify a primitive form of wealth. Proc. Natl. Acad. Sci. 110, 15788–15793 (2013).

4. Vermeer, A. B. L., Boksem, M. A. & Sanfey, A. G. Neural mechanisms underlying context-dependent shifts in risk preferences. NeuroImage 103, 355–363 (2014).

5. Stephens, D. W. Decision ecology: foraging and the ecology of animal decision making. Cogn. Affect. Behav. Neurosci. 8, 475–484 (2008).

6. Tversky, A. & Kahneman, D. Prospect theory: An analysis of decision under risk. Econometrica 47, 263–291 (1979).

7. Ruggeri, K. et al. Replicating patterns of prospect theory for decision under risk. Nat. Hum. Behav. 1–12 (2020).

8. Wakker, P. P. Prospect theory: For risk and ambiguity. (Cambridge university press, 2010).

9. Breiter, H. C., Aharon, I., Kahneman, D., Dale, A. & Shizgal, P. Functional imaging of neural responses to expectancy and experience of monetary gains and losses. Neuron 30, 619–639 (2001).

10. Hsu, M., Bhatt, M., Adolphs, R., Tranel, D. & Camerer, C. F. Neural systems responding to degrees of uncertainty in human decision-making. Science 310, 1680–1683 (2005).

11. Hsu, M., Krajbich, I., Zhao, C. & Camerer, C. F. Neural response to reward anticipation under risk is nonlinear in probabilities. J. Neurosci. 29, 2231–2237 (2009).

12. Jung, W. H., Lee, S., Lerman, C. & Kable, J. W. Amygdala functional and structural connectivity predicts individual risk tolerance. Neuron 98, 394–404. e4 (2018).

13. Kuhnen, C. M. & Knutson, B. The neural basis of financial risk taking. Neuron 47, 763–770 (2005).

14. Huettel, S. A., Stowe, C. J., Gordon, E. M., Warner, B. T. & Platt, M. L. Neural signatures of economic preferences for risk and ambiguity. Neuron 49, 765–775 (2006).

15. Craig, A. D. How do you feel? Interoception: the sense of the physiological condition of the body. Nat. Rev. Neurosci. 3, 655–666 (2002).

16. Craig, A. D. & Craig, A. D. How do you feel--now? The anterior insula and human awareness. Nat. Rev. Neurosci. 10, (2009).

17. Shiv, B., Loewenstein, G. & Bechara, A. The dark side of emotion in decision-making: When individuals with decreased emotional reactions make more advantageous decisions. Cogn. Brain Res. 23, 85–92 (2005).

18. Clark, L. et al. Differential effects of insular and ventromedial prefrontal cortex lesions on risky decision-making. Brain 131, 1311–1322 (2008).

19. Mizuhiki, T., Richmond, B. J. & Shidara, M. Encoding of reward expectation by monkey anterior insular neurons. J. Neurophysiol. 107, 2996–3007 (2012).

20. Kaskan, P. M. et al. Learned value shapes responses to objects in frontal and ventral stream networks in macaque monkeys. Cereb. Cortex 27, 2739–2757 (2017).

21. Tversky, A. & Kahneman, D. The framing of decisions and the psychology of choice. science 211, 453–458 (1981).

22. Bossaerts, P. Risk and risk prediction error signals in anterior insula. Brain Struct. Funct. 214, 645–653 (2010).

23. Preuschoff, K., Quartz, S. R. & Bossaerts, P. Human insula activation reflects risk prediction errors as well as risk. J. Neurosci. 28, 2745–2752 (2008).

24. Tversky, A. & Kahneman, D. Advances in prospect theory: Cumulative representation of uncertainty. J. Risk Uncertain. 5, 297–323 (1992).

25. Farashahi, S., Azab, H., Hayden, B. & Soltani, A. On the flexibility of basic risk attitudes in monkeys. J. Neurosci. 38, 4383–4398 (2018).

26. Stauffer, W. R., Lak, A., Bossaerts, P. & Schultz, W. Economic choices reveal probability distortion in macaque monkeys. J. Neurosci. 35, 3146–3154 (2015).

27. Chen, M. K., Lakshminarayanan, V. & Santos, L. R. How basic are behavioral biases? Evidence from capuchin monkey trading behavior. J. Polit. Econ. 114, 517–537 (2006).

28. Constantinople, C. M., Piet, A. T. & Brody, C. D. An analysis of decision under risk in rats. Curr. Biol. 29, 2066–2074. e5 (2019).

29. So, N.-Y. & Stuphorn, V. Supplementary eye field encodes option and action value for saccades with variable reward. J. Neurophysiol. 104, 2634–2653 (2010).

30. McCoy, A. N. & Platt, M. L. Risk-sensitive neurons in macaque posterior cingulate cortex. Nat. Neurosci. 8, 1220–1227 (2005).

31. Hershey, J. C. & Schoemaker, P. J. Prospect theory’s reflection hypothesis: A critical examination. Organ. Behav. Hum. Perform. 25, 395–418 (1980).

32. Fishburn, P. C. & Kochenberger, G. A. Two-piece von Neumann-Morgenstern utility functions. Decis. Sci. 10, 503–518 (1979).

33. Eisenreich, B. R., Hayden, B. Y. & Zimmermann, J. Macaques are risk-averse in a freely moving foraging task. Sci. Rep. 9, 1–12 (2019).

34. Ogawa, H. Gustatory cortex of primates: anatomy and physiology. Neurosci. Res. 20, 1–13 (1994).

35. Vincis, R., Chen, K., Czarnecki, L., Chen, J. & Fontanini, A. Dynamic representation of taste-related decisions in the gustatory insular cortex of mice. Curr. Biol. (2020).

36. Critchley, H. D. & Garfinkel, S. N. The influence of physiological signals on cognition. Curr. Opin. Behav. Sci. 19, 13–18 (2018).

37. Livneh, Y. et al. Estimation of Current and Future Physiological States in Insular Cortex. Neuron (2020).

38. Nieder, A. & Dehaene, S. Representation of number in the brain. Annu. Rev. Neurosci. 32, 185–208 (2009).

39. Nieder, A. The neuronal code for number. Nat. Rev. Neurosci. 17, 366 (2016).

40. Kutter, E. F., Bostroem, J., Elger, C. E., Mormann, F. & Nieder, A. Single neurons in the human brain encode numbers. Neuron 100, 753–761. e4 (2018).

41. Wang, L., Uhrig, L., Jarraya, B. & Dehaene, S. Representation of numerical and sequential patterns in macaque and human brains. Curr. Biol. 25, 1966–1974 (2015).

42. Tom, S. M., Fox, C. R., Trepel, C. & Poldrack, R. A. The neural basis of loss aversion in decision-making under risk. Science 315, 515–518 (2007).

43. Kahn, I. et al. The role of the amygdala in signaling prospective outcome of choice. Neuron 33, 983–994 (2002).

44. Knutson, B., Fong, G. W., Adams, C. M., Varner, J. L. & Hommer, D. Dissociation of reward anticipation and outcome with event-related fMRI. Neuroreport 12, 3683–3687 (2001).

45. Yacubian, J. et al. Dissociable systems for gain-and loss-related value predictions and errors of prediction in the human brain. J. Neurosci. 26, 9530–9537 (2006).

46. Loewenstein, G. F., Weber, E. U., Hsee, C. K. & Welch, N. Risk as feelings. Psychol. Bull. 127, 267 (2001).

47. Evrard, H. C. The organization of the primate insular cortex. Front. Neuroanat. 13, 43 (2019).

48. Asaad, W. F. & Eskandar, E. N. A flexible software tool for temporally-precise behavioral control in Matlab. J. Neurosci. Methods 174, 245–258 (2008).

49. Seo, H. & Lee, D. Behavioral and neural changes after gains and losses of conditioned reinforcers. J. Neurosci. 29, 3627–3641 (2009).

50. Zandbelt, B. Exgauss: a MATLAB toolbox for fitting the ex-Gaussian distribution to response time data. figshare. (2014).

51. Kahneman, D. & Tversky, A. Prospect theory: An analysis of decision under risk. in HANDBOOK OF THE FUNDAMENTALS OF FINANCIAL DECISION MAKING: Part I 99–127 (World Scientific, 2013).

52. Lattimore, P. K., Baker, J. R. & Witte, A. D. The influence of probability on risky choice: A parametric examination. (1992).

53. Reil, J. C. Die sylvische Grube. Arch Physiol 9, 195–208 (1809).

54. Reveley, C. et al. Three-dimensional digital template atlas of the macaque brain. Cereb. Cortex 27, 4463–4477 (2017).

55. Bakker, R., Tiesinga, P. & Kötter, R. The scalable brain atlas: instant web-based access to public brain atlases and related content. Neuroinformatics 13, 353–366 (2015).

56. Hanes, D. P., Patterson, W. F. & Schall, J. D. Role of frontal eye fields in countermanding saccades: visual, movement, and fixation activity. J. Neurophysiol. 79, 817–834 (1998).

57. Sayer, R. J., Friedlander, M. J. & Redman, S. J. The time course and amplitude of EPSPs evoked at synapses between pairs of CA3/CA1 neurons in the hippocampal slice. J. Neurosci. 10, 826–836 (1990).

